# High salt induces cognitive impairment via the angiotensin II-AT1 and prostaglandin E2-EP1 systems

**DOI:** 10.1101/2022.06.06.495007

**Authors:** Hisayoshi Kubota, Kazuo Kunisawa, Bolati Wulaer, Masaya Hasegawa, Hitomi Kurahashi, Takatoshi Sakata, Hiroyuki Tezuka, Masanori Kugita, Shizuko Nagao, Taku Nagai, Tomoyuki Furuyashiki, Shuh Narumiya, Kuniaki Saito, Toshitaka Nabeshima, Akihiro Mouri

## Abstract

High salt (HS) intake is a known risk factor for hypertension and dementia. Clinical studies have shown that antihypertensive drugs can decrease the incidence of dementia. Accordingly, a strong relationship can be suggested between hypertension and cognitive impairment. It is well-known that angiotensin II (Ang II)-AT1 and prostaglandin E2 (PGE2)-EP1 systems are involved in hypertension and neurotoxicity. However, the involvement of these systems in HS-mediated hypertension and emotional and cognitive impairments remains unclear. Herein, we demonstrated that hypertension and impaired social behavior and object recognition memory following HS intake could be associated with tau hyperphosphorylation, decreased phosphorylation of Ca^2+^/calmodulin-dependent protein kinase II (CaMKII), and postsynaptic density protein 95 (PSD95) expression in the prefrontal cortex and hippocampus of mice. These changes were blocked by pharmacological treatment with losartan, an Ang II receptor blocker (ARB), or EP1 gene knockout. Our findings suggest that Ang II-AT1 and PGE2-EP1 systems could be novel therapeutic targets for hypertension-induced dementia.

## Introduction

Dementia is characterized by cognitive impairment, including memory loss, poor judgment, and abnormalities in social behavior (Arvanitakis, Shah et al., 2019). Approximately 50 million people worldwide are currently living with dementia, and the number of patients is predicted to increase to more than 130 million by 2050 (Disease, Injury et al., 2017, Frisoni, Altomare et al., 2022). As mixed dementia, defined as the coexistence of Alzheimer’s disease (AD) and vascular dementia (VaD), is considered the most common type of dementia, vascular disorders could be implicated as potential underlying causes of dementia (Iadecola, 2013). Longitudinal studies have indicated that lifestyle-related diseases, including hypertension, dyslipidemia, and diabetes, are associated with the subsequent development of dementia (Kivipelto, Mangialasche et al., 2018). Given that midlife blood pressure levels are positively associated with late-life onset VaD, hypertension has been implicated as a significant risk factor for dementia (Ninomiya, Ohara et al., 2011). Clinical studies have shown that antihypertensive drugs can decrease the incidence of dementia (Ihara & Saito, 2020). These findings suggest that cognitive impairment can be prevented by controlling blood pressure.

Numerous preclinical studies have reported that high salt (HS) intake induces vascular lesions and cognitive impairment. Reportedly, hypertensive rats fed an HS diet showed cognitive impairment with abnormalities in synaptic plasticity (Guo, Wei et al., 2017). It is well-known that hyperphosphorylation of the microtubule-associated protein tau increases neuronal loss and cognitive impairment in AD and dementia (Brunden, Trojanowski et al., 2009). A recent study has reported that tau hyperphosphorylation mediates HS-induced cognitive impairment (Faraco, Hochrainer et al., 2019). HS intake plays a critical role in dementia pathology. However, the detailed mechanism through which HS intake induces cognitive impairment via tau hyperphosphorylation remains poorly understood.

The renin-angiotensin system (RAS) is a hormone system that regulates body fluid, electrolyte balance, and blood pressure (Sparks, Crowley et al., 2014). The RAS plays a critical role in the development of vascular disorders, including hypertension (Sparks et al., 2014). In the RAS, angiotensin (Ang) II is synthesized by angiotensin-converting enzyme (ACE) and acts on two receptors, angiotensin II type 1 (AT1) and type 2 (AT2) receptors. AT1 receptor activation results in elevated blood pressure, oxidative stress, neurotoxicity, and cognitive impairment (Cosarderelioglu, Nidadavolu et al., 2020, Villapol & Saavedra, 2015). Conversely, the AT2 receptor has a beneficial role on cerebral blood flow, neuronal plasticity, and learning and memory (Cosarderelioglu et al., 2020, Villapol & Saavedra, 2015). In postmortem studies, elevated ACE-1 activity (Miners, Ashby et al., 2008) and high levels of Ang II (Kehoe, Wong et al., 2016) and AT1 receptors (Cosarderelioglu, Nidadavolu et al., 2021) were documented in the brains of patients with AD. Intracerebroventricular injection of Ang II can impair spatial learning and memory in rats with increased levels of β-amyloid (Aβ) and phosphorylated tau via the AT1 receptor (Tian, Zhu et al., 2012, Zhu, Shi et al., 2011). ACE inhibitors and AT1 receptor blockers (ARB) reportedly induce beneficial effects on neurodegeneration and cognitive impairment (AbdAlla, Langer et al., 2013, Dong, Kataoka et al., 2011, Takeda, Sato et al., 2009, Tian et al., 2012). These reports suggest that the Ang II-AT1 system may be associated with cognitive impairment following HS intake.

Prostaglandin E2 (PGE2) is produced by sequential activation of cyclooxygenase (COX)-1,2 enzymes, microsomal prostaglandin E synthases (mPGES)-1,2, and cytosolic PGES (cPGES) in the arachidonic acid cascade and exhibits various effects on blood pressure and the central nervous system (CNS) via four types receptors (EP1-EP4) (Breyer & Breyer, 2001, Mohan, Ahmad et al., 2012). EP1 is expressed in diverse tissues, but predominantly in the kidney, mediating the effect of PGE2 on vasoconstriction in the peripheral vasculature (Breyer & Breyer, 2001). In the CNS, EP1 is known to participate in neurotoxicity in cerebral ischemia and AD (Kawano, Anrather et al., 2006, Zhen, Kim et al., 2012). It has been reported that the Ang II-AT1 system regulates COX-2 expression and PGE2 production in the kidney (Quadri, Culver et al., 2016). Interestingly, Ang II-induced hypertension was attenuated by pharmacological and genetic inhibition of EP1 (Guan, Zhang et al., 2007). The PGE2-EP1 system is involved in Ang II-induced reactive oxygen species accumulation in the subcortical region of the brain (Cao, Peterson et al., 2012). The interaction between Ang II-AT1 and PGE2-EP1 systems might play an important role in the development of hypertension and dementia.

To clarify this hypothesis, we examined whether Ang II-AT1 and PGE2-EP1 systems are associated with hypertension and emotional and cognitive impairments following HS intake. Our findings provide novel therapeutic targets for dementia accompanied by HS-induced hypertension.

## Results

### Hypertension and impairments in social behavior and object recognition memory after HS intake

To establish an animal model of hypertension, mice were provided an HS solution (2% NaCl drinking water) instead of water for 12 weeks (Fig 1A). The mice were monitored weekly to determine body weight, water consumption, heart rate, and systolic blood pressure. Compared with normal water intake in the control group, HS intake did not alter body weight (Fig 1B) but increased water consumption in mice (Fig 1C). Systolic blood pressure was elevated 6 weeks after HS intake (Fig 1E), although no significant difference in heart rate was detected between the groups (Fig 1D). No significant differences in the serum biochemical profiles (TP, Alb, T-Bil, ALP, AST, ALT, LD, γ-GT, T-cho, HDL-cho, TG, UA, Glu, BUN, and Cre) were noted between groups, suggesting that HS intake did not impair hepatic or renal functions (Table EV1). To determine the effect of HS intake on electrolyte homeostasis, Na^+^, K^+^, and Cl^−^ levels in the serum and urine of mice were measured. HS intake did not alter serum levels of Na^+^, K^+^, or Cl^−^ (Fig EV1A-C). However, urine levels of Na^+^ and Cl^−^ but not K^+^ were markedly increased by HS intake (Fig EV1D-F). These data suggested that HS-treated mice excreted excessive Na^+^ and Cl^−^ into the urine to control electrolyte homeostasis.

**Figure 1.**
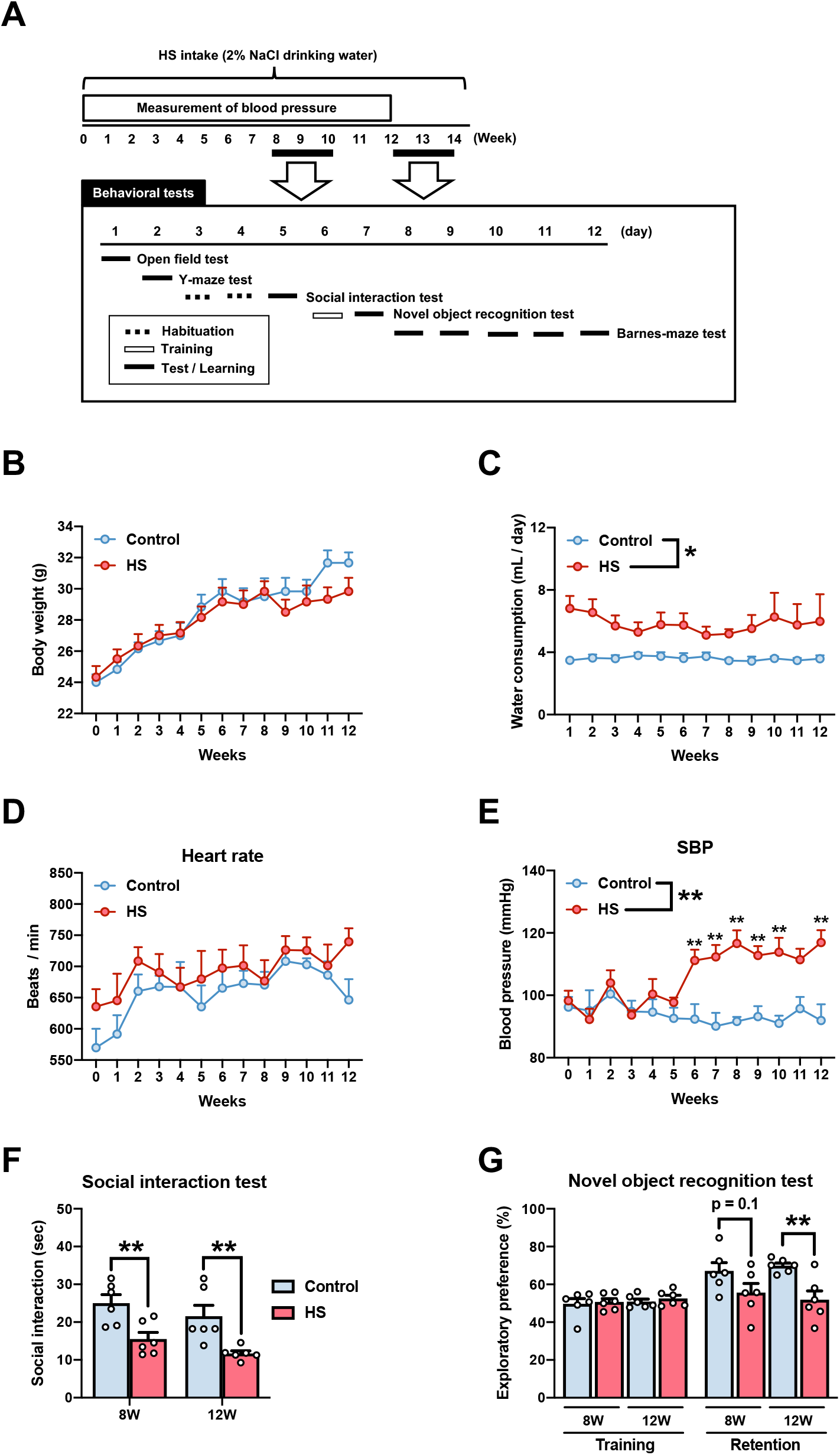
HS intake induces hypertension and impairments in social behavior and object recognition memory. **A** Protocol for the HS intake model. **B-E** (B) Body weight, (C) water consumption, (D) heart rate, and (E) systolic blood pressure of experimental animals were monitored once weekly for 12 weeks. (C) Water consumption was determined based on an average of three home cages. (D) Heart rate and (E) systolic blood pressure were measured by the tail-cuff system. **F, G** The mice were subjected to behavioral tests at 8 and 12 weeks after HS intake. (F) Social behavior in the social interaction test. The mice were placed in an apparatus with an unfamiliar mouse, and their social interaction was measured for 10 min. (G) Object recognition memory in the novel object recognition test. The retention session was performed 24 h after the training session. Exploratory preference was measured during a 10 min session. Data information: Each column represents the mean ± standard error of the mean (SEM) (n=6). *p < 0.05, **p < 0.01 versus control. (B)-(E) Two-way ANOVA followed by Tukey’s multiple comparison test: (B) *F* _Weeks (12,120)_ = 42.96, *p* < 0.01; *F* _HS (1,10)_ = 0.23, *p* = 0.64; *F* _Weeks × HS (12,120)_ = 2.15, *p* < 0.05, (C) *F* _Weeks (11,44)_ = 0.61, *p* = 0.81; *F* _HS (1,4)_ = 7.72, *p* < 0.05; *F* _Weeks × HS (11,44)_ = 0.73, *p* = 0.70, (D) *F* _Weeks (12,120)_ = 4.00, *p* < 0.01; *F* _HS (1,10)_ = 1.74, *p* = 0.22; *F* _Weeks × HS (12,120)_ = 0.61, *p* = 0.83, (E) *F* _Weeks (12,120)_ = 2.13, *p* < 0.05; *F* _HS (1,10)_ = 73.2, *p* < 0.01; *F* _Weeks × HS (12,120)_ = 3.82, *p* < 0.01. (F), (G) Student’s t-test: (F) 8 weeks, t(10) = 3.34, p < 0.01; 12 weeks, t(10) = 3.30, p < 0.01, (G) Training: 8 weeks, t(10) = 0.34, p = 0.75; 12 weeks, t(10) = 0.82, p = 0.43; Retention: 8 weeks, t(10) = 1.76, p = 0.11; 12 weeks, t(10) = 3.66, p < 0.01. HS, high salt; SBP, systolic blood pressure.

Overall, these results demonstrated that HS intake induces hypertension without a significant influence on hepatic and renal functions.

Previous studies have shown that hypertension can be associated with emotional and cognitive impairments and significantly increases the risk of dementia (Cohen, Edmondson et al., 2015, Ungvari, Toth et al., 2021). To examine whether HS intake alters emotional and cognitive functions, mice underwent several types of behavioral tests 8 and 12 weeks after HS intake (Fig 1A). In the social interaction test, the duration of social interaction significantly decreased 8 and 12 weeks after HS intake (Fig 1F). In the novel object recognition test, no significant differences were documented in exploratory preference (Fig 1G) and the total time taken to explore the two objects (Fig EV2E) during the training session. During the retention session, exploratory preference for the novel object decreased 12 weeks after HS intake (Fig 1G); however, the total time taken to explore the two objects was unaltered (Fig EV2E). To further evaluate emotional and cognitive functions, the mice were subjected to the open field, Y-maze, and Barnes-maze paradigms. However, there were no significant differences between groups in terms of time spent in each zone (Fig EV2A) and locomotor activity (Fig EV2B) in the open field test, spontaneous alternation (Fig EV2C) and total arm entries (Fig EV2D) in the Y-maze test, and latency to enter the target escape box (Fig EV2F) in the Barnes maze test. These results demonstrated that impairments in social behavior and object recognition memory were observed 8 and 12 weeks after HS intake, respectively.

Accordingly, these findings suggested that social behavior is more sensitive to HS intake than recognition memory.

### Tau hyperphosphorylation and reduction of synapse-related proteins in the PFC and HIP after HS intake

The connectivity of the neuronal network, including the PFC and HIP, is strongly implicated in social behavior and object recognition memory (Ko, 2017, Sorooshyari, Sheng et al., 2020). Previous studies have reported that an HS diet induces tau hyperphosphorylation and downregulates synapse-related proteins in the rodent brain (Faraco et al., 2019, Guo et al., 2017). We next confirmed whether HS intake increased tau phosphorylation and impaired synaptic function in the PFC and HIP. The phosphorylation-specific tau antibody (AT8), which detects Ser^202^ and Thr^205^ sites, is widely employed to evaluate tau pathology in AD brains (Braak, Alafuzoff et al., 2006, Lee, Goedert et al., 2001). We detected high levels of phosphorylated tau (AT8) immunoreactivity in the PFC of HS-treated mice (Fig 2A). Tau is phosphorylated by various serine and threonine residues (Simic, Babic Leko et al., 2016). We examined tau phosphorylation using not only AT8 but also other antibodies to detect phosphorylation at Ser^202^, Ser^396^, Ser^404^ and Ser^416^ sites by western blotting (Fig 2B and C). Twelve weeks after HS intake, tau hyperphosphorylation was increased at the AT8, Ser^396^, and Ser^404^ sites, but not at Ser^202^ and Ser^416^ sites in the PFC (Fig 2B). In the HIP, tau hyperphosphorylation was increased at the AT8 and Ser^404^ sites12 weeks after HS intake, but not at the Ser^202^, Ser^396^, and Ser^416^ sites (Fig 2B). Time-course analysis revealed tau hyperphosphorylation at the AT8 site 8 weeks after HS intake (Fig 2C).

**Figure 2.**
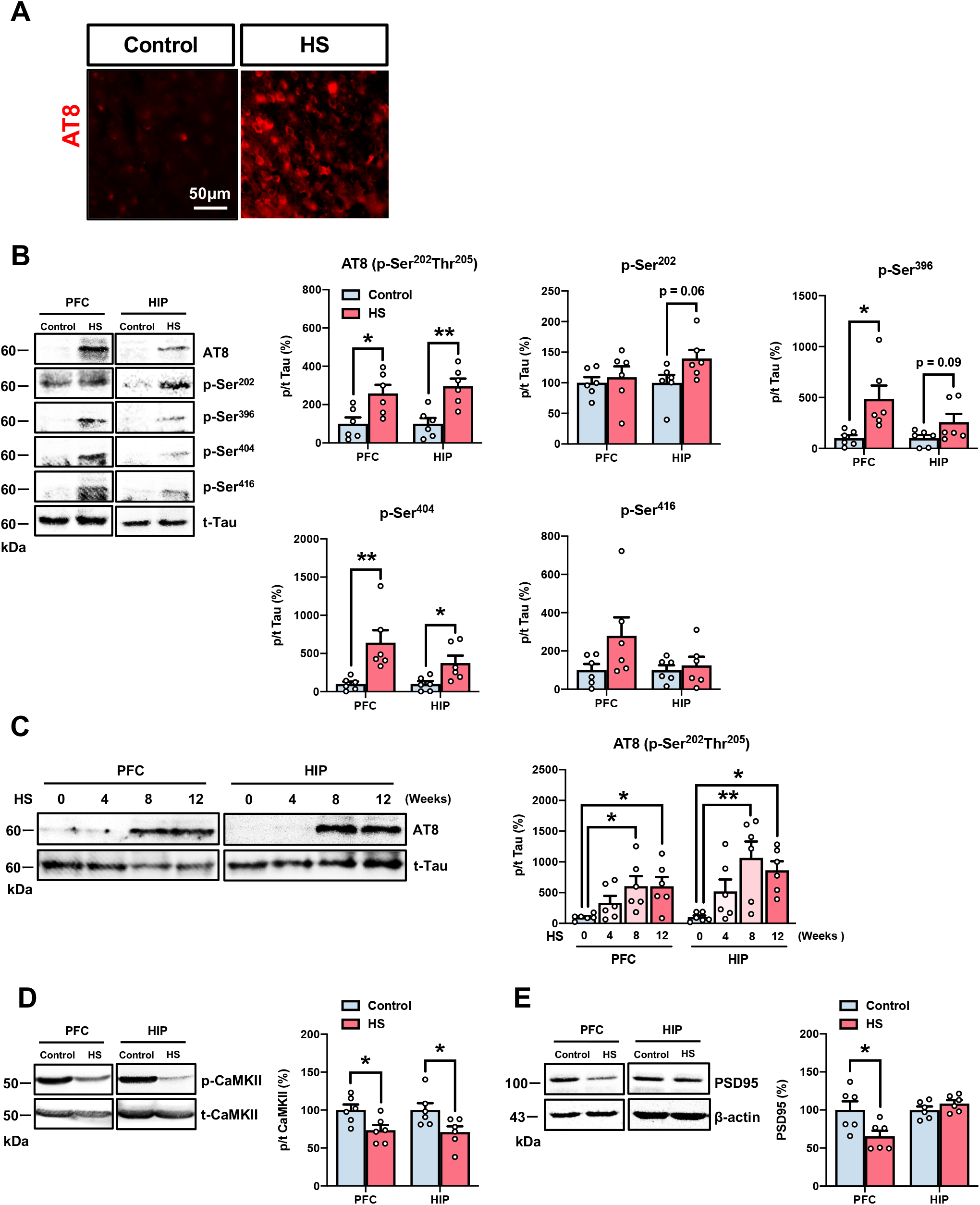
Tau hyperphosphorylation and reduction of synapse-related proteins in the prefrontal cortex and hippocampus after HS intake. **A** Immunostaining of AT8 (red) was performed using phosphorylation-specific tau antibody in the prefrontal cortex of mice 12 weeks after HS intake. Scale bar= 50 μm. **B-E** Expression levels of phosphorylated tau and synapse-related proteins were measured by western blotting in the prefrontal cortex and hippocampus of mice 12 weeks after HS intake. (C) Time-course analysis for AT8 in the prefrontal cortex and hippocampus of mice 4, 8, and 12 weeks after the HS intake was performed by western blotting. The phosphorylation ratio of (B), (C) AT8, Ser^202^, Ser^396^, Ser^404^, Ser^416^, and (D) CaMKII and protein levels of (E) PSD95 in the prefrontal cortex and hippocampus were calculated. Loaded protein was normalized to actin. The phosphorylation ratio was calculated as tau phosphorylation versus t-Tau expression and CaMKII phosphorylation versus t-CaMKII expression. Data information: Each column represents the mean ± standard error of the mean (SEM) (n =6). *p < 0.05, **p < 0.01 versus control. (B), (D), (E) Student’s t-test: (B) AT8: PFC, t(10) = 2.86, p < 0.05; HIP, t(10) = 3.99, p < 0.01, Ser^202^: PFC, t(10) = 0.45, p = 0.66; HIP, t(10) = 2.08, p = 0.06, Ser^396^: PFC, t(10) = 2.85, p < 0.05; HIP, t(10) = 1.83, p = 0.09, Ser^404^: PFC, t(10) = 3.26, p < 0.01; HIP, t(10) = 2.62, p < 0.05, Ser^416^: PFC, t(10) = 1.75, p = 0.11; HIP, t(10) = 0.47, p = 0.65, (D) PFC, t(10) = 2.68, p < 0.05; HIP, t(10) = 2.44, p < 0.05, (E) PFC, t(10) = 2.56, p < 0.05; HIP, t(10) = 1.33, p = 0.21. (C) One-way ANOVA followed by Tukey’s multiple comparison test: PFC, *F* _(3,20)_ = 3.80, *p* < 0.05; HIP, *F* _(3,20)_ = 5.47, *p* < 0.01. HS, high salt; PFC, prefrontal cortex; HIP, hippocampus; AT8, phosphorylated tau at Ser^202^/Thr^205^ site; CaMKII, Ca^2+^/calmodulin-dependent protein kinase II; PSD95, postsynaptic density protein 95.

Cyclin-dependent kinase 5 (CDK5) is a major kinase involved in tau phosphorylation. The activation of the calcium-dependent protease calpain cleaves p35 to p25, which can activate CDK5, resulting in tau phosphorylation (Castro-Alvarez, Uribe-Arias et al., 2014). HS intake decreased calpain expression in the PFC but not in the HIP (Fig EV3A), and did not affect the expression of p25 in the PFC or HIP (Fig EV3B).

Accumulation of pathological tau induces synaptic impairment (Wu, Zhang et al., 2021, Ye, Yin et al., 2020, Yin, Gao et al., 2016). We next investigated the expression of synapse-related proteins, phosphorylated Ca^2+^/calmodulin-dependent protein kinase II (CaMKII), and postsynaptic density protein 95 (PSD95), by western blotting. Twelve weeks after HS intake, the expression of phosphorylated CaMKII was decreased in the PFC and HIP (Fig 2D). PSD95 expression decreased only in the PFC (Fig 2E).

Furthermore, several clinical studies have revealed impairment of monoaminergic systems in the postmortem brains of patients with AD (Simic, Babic Leko et al., 2017). However, 12 weeks after HS intake, no differences in the content of monoamines and their metabolites were detected in the PFC, nucleus accumbens, striatum, HIP, and amygdala (Table EV2).

Taken together, our results suggested that HS intake induced tau hyperphosphorylation and reduction of synapse-related proteins without altering monoaminergic systems.

### The degenerated neuronal morphology of PrL and CA1 neurons after HS intake

Given that HS intake induced tau hyperphosphorylation and reduced synapse-related proteins, we performed Golgi stating to examine the neuronal morphology of the PrL subregion in the PFC (Fig 3A) and hippocampal CA1 subregions (Fig 3E). Twelve weeks after HS intake, treated mice showed a shortened total dendritic length in the PrL (Fig 3B). On assessing the effect of HS intake on dendritic branching by Sholl analysis, HS-treated mice showed shortened dendritic length (Fig 3C) and reduced intersections (Fig 3D) per Sholl-segment, particularly those 20–300 μm from the soma in the PrL. Similar results were observed in the CA1 region of HS-treated mice (total dendritic length, Fig 3F; dendritic length, Fig 3G; intersections, Fig 3H).

**Figure 3.**
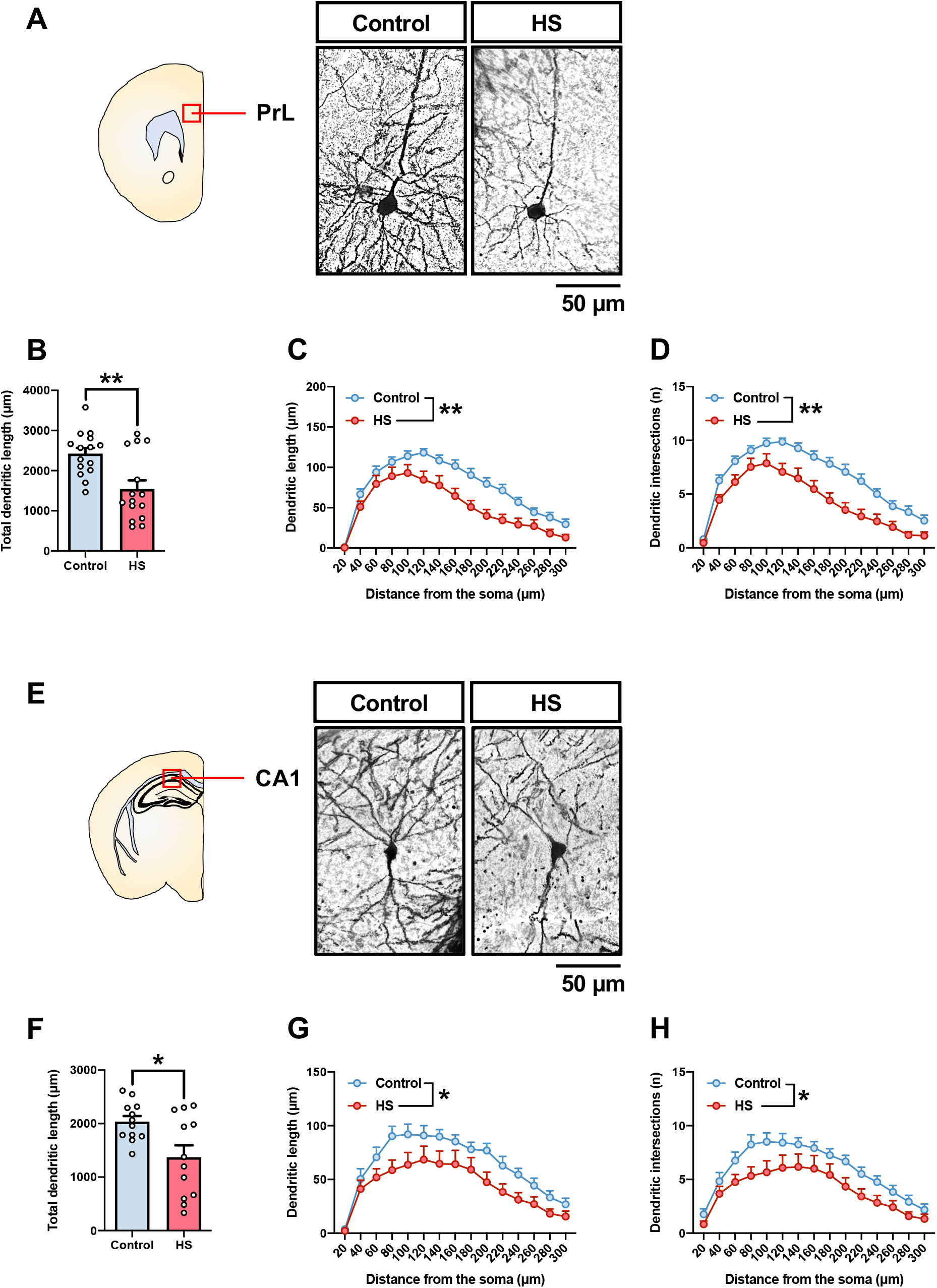
Degenerated neuronal morphology of PrL and CA1 neurons after HS intake. **A-H** The morphology of PrL and CA1 neurons in mice 12 weeks after HS intake was analyzed by Golgi staining. (A) Representative images showing PrL neurons in control and HS-treated mice. Quantification of (B) total dendritic length, (C) dendritic length, and (D) dendritic intersection in PrL neurons. (E) Representative images showing CA1 neurons in control and HS-treated mice. Quantification of (F) total dendritic length, (G) dendritic length, and (H) dendritic intersections in CA1 neurons. Scale bars= 50 µm. Data information: Each column represents the mean ± standard error of the mean (SEM) (n = 12-15 dendrites from 4-5 mice). *p < 0.05, **p < 0.01 versus control. (B), (F) Student’s t-test: (B) t(28) = 3.43, p < 0.01, (F) t(22) = 2.72, p < 0.05. (C), (D), (G), (H) Two-way ANOVA followed by Tukey’s multiple comparison test: (C) *F* _Distance from the soma (14,392)_ = 74.28, *p* < 0.01; *F* _HS (1,28)_ = 9.77, *p* < 0.01; *F* _Distance from the soma × HS (14,392)_ = 2.39, *p* < 0.01, (D) *F* _Distance from the soma (14,392)_ = 78.39, *p* < 0.01; *F* _HS (1,28)_ = 13.63, *p* < 0.01; *F* _Distance from the soma × HS (14,392)_ = 2.18, *p* < 0.01, (G) *F* _Distance from the soma (14,308)_ = 30.20, *p* < 0.01; *F* _HS (1,22)_ = 6.01, *p* < 0.05; *F* _Distance from the soma × HS (14,308)_ = 0.88, *p* = 0.58, (H) *F* _Distance from the soma (14,308)_ = 27.27, *p* < 0.01; *F* _HS (1,22)_ = 6.05, *p* < 0.05; *F* _Distance from the soma × HS (14,308)_ = 0.61, *p* = 0.85. HS, high salt; PrL, prelimbic region of the prefrontal cortex; CA1, CA1 region of the hippocampus.

These results suggested that HS intake leads to dendritic degeneration in the PFC and HIP.

### Changes in the RAS-related gene expression after HS intake

Ang II is generated by the RAS and plays a critical role in blood pressure homeostasis via the AT1 receptor (Oparil, Acelajado et al., 2018). Our results showed that HS intake induced hypertension (Fig 1E). To investigate whether HS intake alters RAS, the levels of mRNA related to RAS were measured in the kidney, PFC, and HIP of mice. Twelve weeks after HS intake, the mRNA levels of angiotensinogen (AGT) and ACE were increased in the kidney and PFC, but renin in the HIP and AT1 in the PFC were decreased (Fig 4A-E).

**Figure 4.**
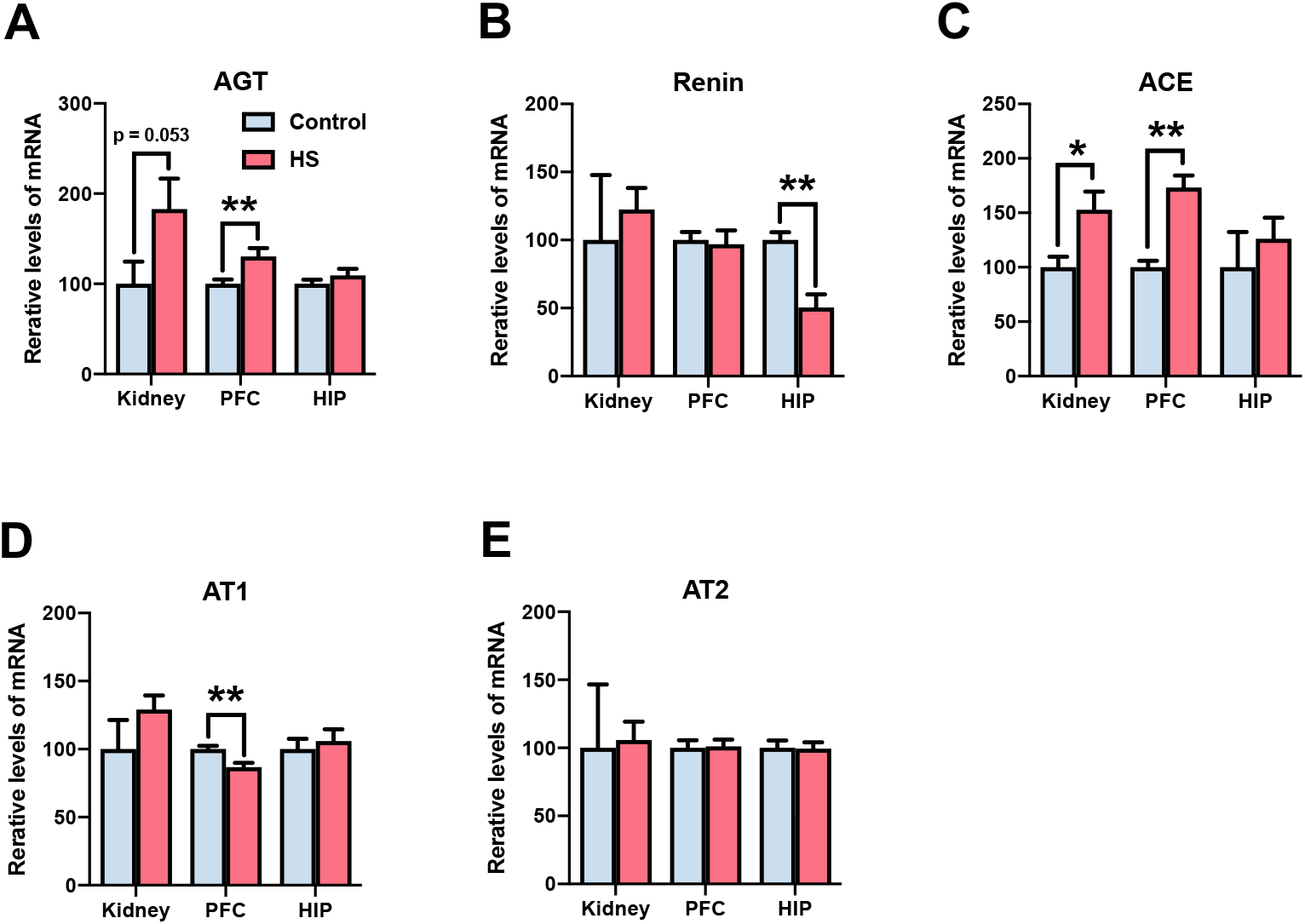
The change in the RAS-related gene expressions after HS intake. **A-E** The mRNA expression of (A) AGT, (B) renin, (C) ACE, (D) AT1, and (E) AT2 was measured by real-time reverse transcription-PCR in the kidneys, prefrontal cortex, and hippocampus of mice 12 weeks after HS intake. Data information: Each column represents the mean ± standard error of the mean (SEM) (n =6-8). *p < 0.05, **p < 0.01 versus control. (A)-(E) Student’s t-test: (A) Kidney, t(10) = 2.19, p = 0.053; PFC, t(14) = 3.18, p < 0.05; HIP, t(14) = 1.12, p = 0.28, (B) Kidney, t(10) = 0.50, p = 0.63; PFC, t(14) = 0.27, p = 0.79; HIP, t(14) = 3.71, p < 0.05, (C) Kidney, t(10) = 3.07, p < 0.05; PFC, t(14) = 6.48, p < 0.01; HIP, t(14) = 0.73, p = 0.47, (D) Kidney, t(10) = 1.22, p = 0.25; PFC, t(14) = 3.33, p < 0.01; HIP, t(14) = 0.53, p = 0.61, (E) Kidney, t(10) = 0.14, p = 0.89; PFC, t(14) = 0.15, p = 0.88; HIP, t(14) = 0.09, p = 0.93. HS, high salt; PFC, prefrontal cortex; HIP, hippocampus; AGT, angiotensinogen; ACE, angiotensin-converting enzyme; AT1, angiotensin II type 1 receptor; AT2, angiotensin II type 2 receptor.

These results suggested that HS intake activated the RAS.

### Losartan blocks hypertension, behavioral impairments, and tau hyperphosphorylation after HS intake

We next investigated the involvement of the RAS in hypertension and behavioral impairments following HS intake using losartan, a non-blood-brain barrier (BBB)-crossing ARB. Given that hypertension was observed 6 weeks after HS intake (Fig 1E), mice were administered losartan (i.p.) from 6 weeks after HS intake to the end of the behavioral testing period, and systolic blood pressure was monitored. Subsequently, locomotor activity, sociability, and object recognition memory of mice were examined (Fig 5A). Losartan blocked hypertension after HS intake (Fig 5B). Furthermore, losartan blocked the impaired social behavior and object recognition memory after HS intake (Fig 5C and D), but not locomotor activity (Fig EV4A) or the total time taken to explore the two objects (Fig EV4C).

**Figure 5.**
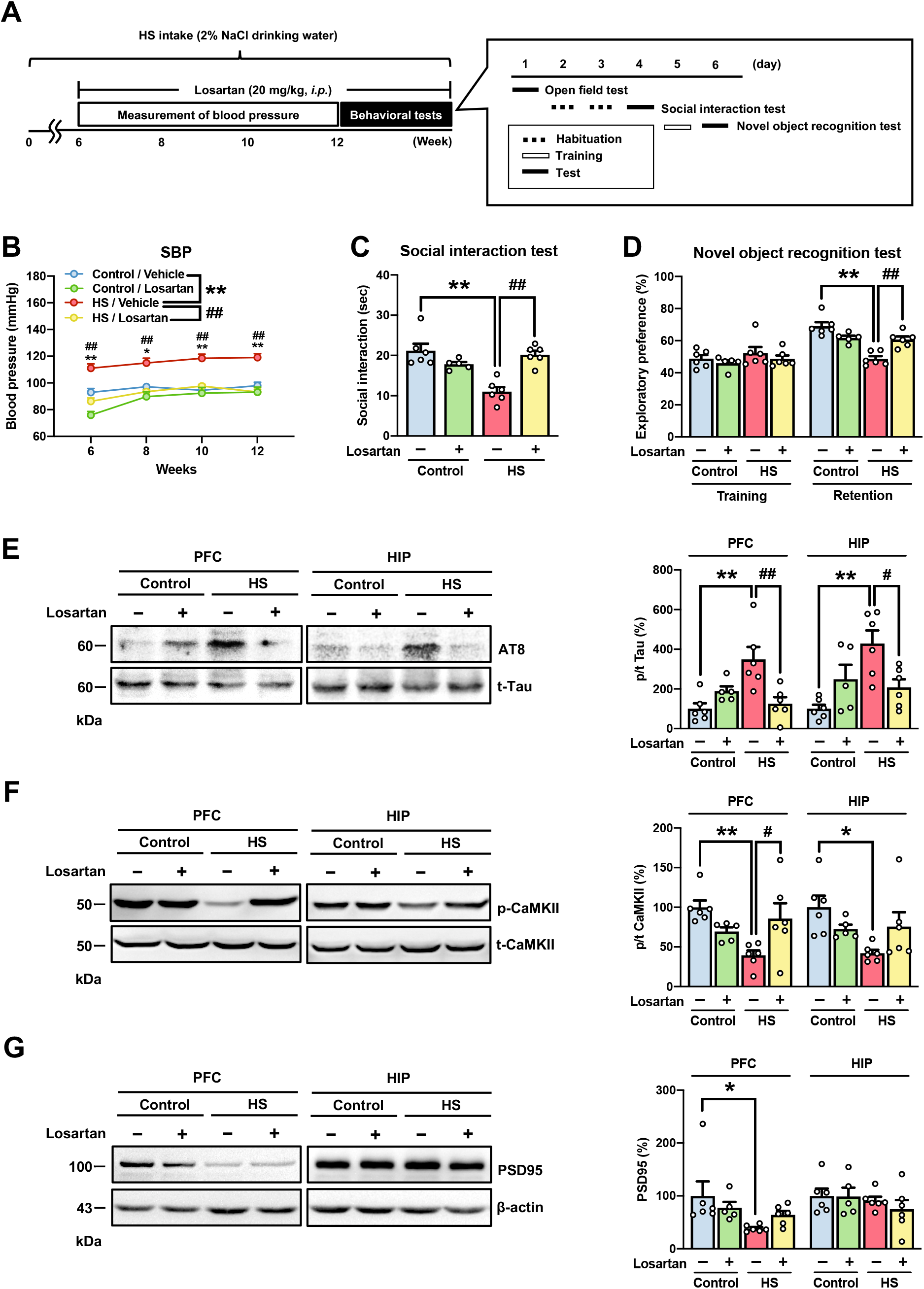
Losartan blocks HS-induced hypertension, behavioral impairments, and tau hyperphosphorylation. **A** Experimental design for evaluating the effects of losartan in the HS intake model. **B** The systolic blood pressure of mice was monitored using the tail-cuff system every two weeks from 6 to 12 weeks. **C, D** The mice were subjected to behavioral tests 12 weeks after HS intake. (C) Social behavior in the social interaction test. Mice were placed in an apparatus with an unfamiliar mouse, and their social interaction was measured for 10 min. (D) Object recognition memory in the novel object recognition test. The retention session was performed 24 h after the training session. Exploratory preference was measured during a 10 min session. **E-G** Expression levels of phosphorylated tau and synapse-related proteins in the prefrontal cortex and hippocampus of mice 12 weeks after the HS intake were measured by western blotting. The phosphorylation ratio of (E) AT8 and (F) CaMKII and the protein levels of (G) PSD95 in the prefrontal cortex and hippocampus were calculated. Loaded protein was normalized to actin. The phosphorylation ratio was calculated as tau phosphorylation versus t-Tau expression and CaMKII phosphorylation versus t-CaMKII expression. Data information: Each column represents the mean ± standard error of the mean (SEM) (n =5-6). *p < 0.05, **p < 0.01 versus control/vehicle, #p < 0.05, ##p < 0.01 versus HS/ vehicle. (B)-(G) Two-way ANOVA followed by Tukey’s multiple comparison test: (B) *F* _Weeks (2.91,55.32)_ = 13.97, *p* < 0.01; *F* _Group (3,19)_ = 40.80, *p* < 0.01; *F* _Weeks × Group (9,57)_ = 1.67, *p* = 0.12, (C) *F* _HS (1,19)_ = 10.23, *p* < 0.01; *F* _Losartan (1,19)_ = 5.64, *p* < 0.05; *F* _HS × Losartan (1,19)_ = 26.53, *p* < 0.01, (D) Training, *F* _HS (1,19)_ = 1.40, *p* = 0.25; *F* _Losartan (1,19)_ = 1.48, *p* = 0.24; *F* _HS × Losartan (1,19)_ = 0.02, *p* = 0.88; Retention, *F* _HS (1,19)_ = 29.28, *p* < 0.01; *F* _Losartan (1,19)_ = 1.47, *p* = 0.24; *F* _HS × Losartan (1,19)_ = 24.05, *p* < 0.01, (E) PFC, *F* _HS (1,19)_ = 5.13, *p* < 0.05; *F* _Losartan (1,19)_ = 2.68, *p* = 0.12; *F* _HS × Losartan (1,19)_ = 14.39, *p* < 0.01; HIP, *F* _HS (1,19)_ = 7.57, *p* < 0.05; *F* _Losartan (1,19)_ = 0.47, *p* = 0.50; *F* _HS × Losartan (1,19)_ = 12.50, *p* < 0.01, (F) PFC, *F* _HS (1,19)_ = 3.51, *p* = 0.08; *F* _Losartan (1,19)_ = 0.45, *p* = 0.51; *F* _HS × Losartan (1,19)_ = 10.58, *p* < 0.01; HIP, *F* _HS (1,19)_ = 4.72, *p* < 0.05; *F* _Losartan (1,19)_ = 0.06, *p* = 0.81; *F* _HS × Losartan (1,19)_ = 5.79, *p* < 0.05, (G) PFC, *F* _HS (1,19)_ = 5.71, *p* < 0.05; *F* _Losartan (1,19)_ = 0.01, *p* = 0.91; *F* _HS × Losartan (1,19)_ = 2.33, *p* = 0.14; HIP, *F* _HS (1,19)_ = 1.36, *p* = 0.26; *F* _Losartan (1,19)_ = 0.37, *p* = 0.55; *F* _HS × Losartan (1,19)_ = 0.29, *p* = 0.60. HS: high salt, PFC, prefrontal cortex; HIP, hippocampus; Losartan, a non-BBB-crossing Ang II receptor blocker; SBP, systolic blood pressure; AT8, phosphorylated tau at Ser^202^/Thr^205^ site; CaMKII, Ca^2+^/calmodulin-dependent protein kinase II; PSD95, postsynaptic density protein 95.

As losartan could block the abnormal behaviors after HS intake, we investigated whether losartan blocked tau hyperphosphorylation and reduced synapse-related proteins in the PFC and HIP of mice by western blotting. We observed that losartan blocked HS intake-induced tau hyperphosphorylation (Fig 5E) and decreased CaMKII phosphorylation without impacting PSD95 expression in the PFC and HIP (Fig 5F and G).

These results suggested that hypertension associated with activation of the Ang II-AT1 system after HS intake could lead to behavioral impairments through tau hyperphosphorylation and reduction in synapse-related proteins in mice.

### Activation of the PGE2-EP1 system in the kidney and brain after HS intake

As shown in Fig 4A and C, we detected elevated mRNA levels of AGT and ACE in the kidney and PFC after HS intake, thus suggesting the activation of the Ang II-AT1 system. RAS-generated Ang II activates the PGE2-EP1 system via AT1 receptor (Quadri et al., 2016, Young & Davisson, 2015). Furthermore, PGE2 mediates various effects on blood pressure and the CNS via four types of PGE2 receptors (EP1-EP4) (Breyer & Breyer, 2000, Milatovic, Montine et al., 2011, Narumiya, Sugimoto et al., 1999). Next, we investigated the interaction between the Ang II-AT1 and PGE2-EP1 systems in HS-treated mice. Twelve weeks after HS intake, the plasma content of PGE2 increased, but losartan blocked this increase in PGE2 levels (Fig 6A). Second, we examined whether HS intake increases mRNA levels of PGE2 synthesizing enzymes and EP receptor subtypes in the kidney, PFC, and HIP. After HS intake, mRNA levels of COX-1, COX-2, and mPGES-1, but not mPGES-2, were increased in the kidney, but not in the PFC and HIP (Fig 6B-E). Additionally, HS intake increased the mRNA levels of EP1, EP3, and EP4 in the kidneys and EP1 in the PFC (Fig 6F-I).

**Figure 6.**
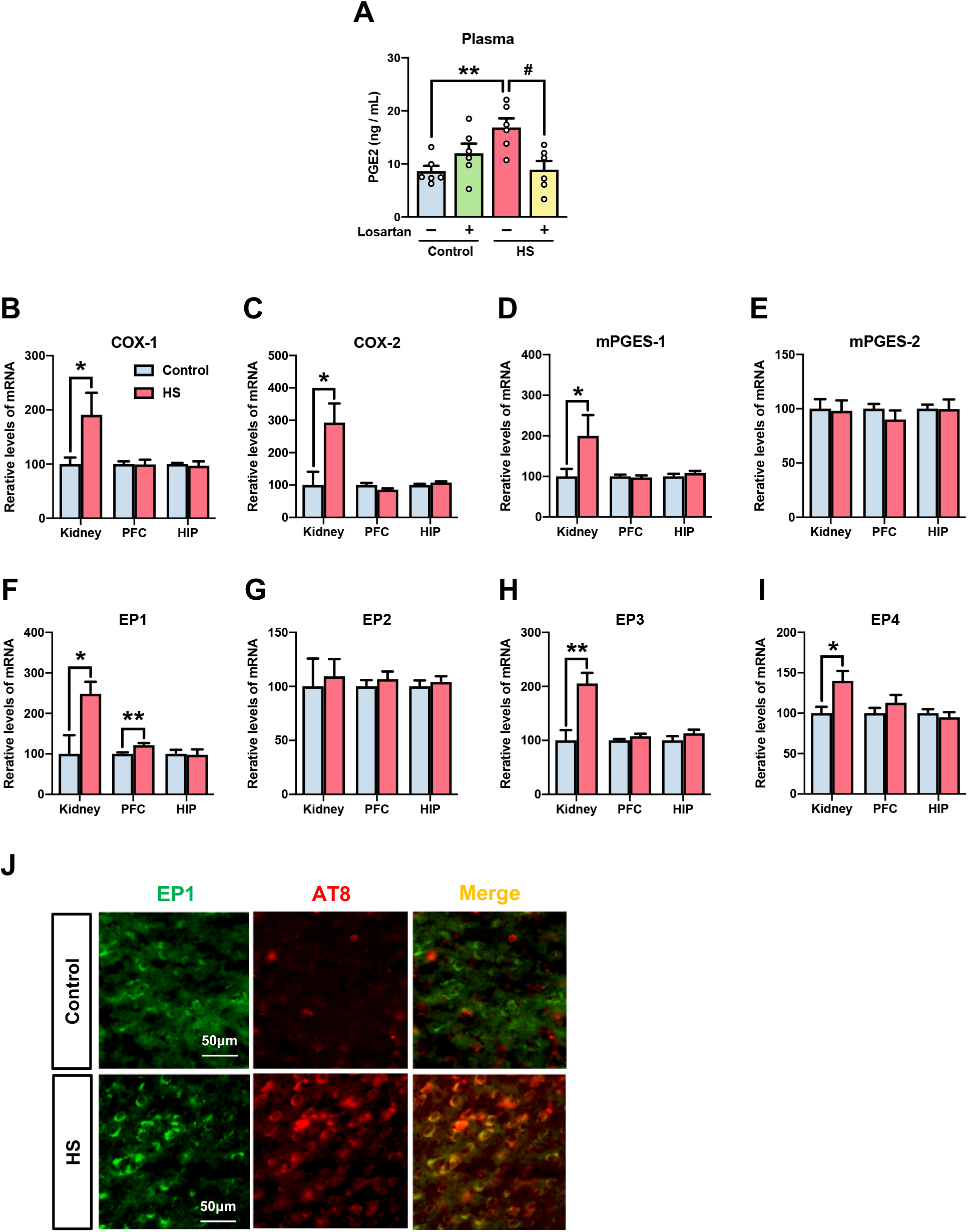
Activation of the PGE2-EP1 system in the kidney and brain after HS intake. **A** Content of PGE2 was measured by ELISA in the plasma of losartan-treated mice 12 weeks after HS intake. **B-I** mRNA expression levels of (B) COX-1, (C) COX-2, (D) mPGES-1, (E) mPGES-2, (F) EP1, (G) EP2, (H) EP3, and (I) EP4 were measured in the kidney, prefrontal cortex, and hippocampus of mice 12 weeks after the HS intake using quantitative reverse transcription-PCR. **J** Co-immunostaining of AT8 and EP1 was performed by using a phosphorylation-specific tau antibody in the prefrontal cortex of mice 12 weeks after the HS intake. Scale bar= 50 μm. Data information: Each column represents the mean ± standard error of the mean (SEM) (n =6-8). *p < 0.05, **p < 0.01 versus control, #p < 0.05 versus HS/ vehicle. (A) Two-way ANOVA followed by Tukey’s multiple comparison test: *F* _HS (1,20)_ = 2.66, *p* = 0.12; *F* _Losartan (1,20)_ = 2.08, *p* = 0.17; *F* _HS × Losartan (1,20)_ = 12.71, *p* < 0.01. (B)-(I) Student’s t-test: (B) Kidney, t(10) = 2.86, p < 0.05; PFC, t(14) = 0.05, p = 0.96; HIP, t(14) = 0.38, p = 0.71, (C) Kidney, t(10) = 2.76, p < 0.05; PFC, t(14) = 1.93, p = 0.07; HIP, t(14) = 1.42, p = 0.18, (D) Kidney, t(10) = 2.45, p < 0.05; PFC, t(14) = 0.37, p = 0.72; HIP, t(14) = 1.05, p = 0.31, (E) Kidney, t(10) = 0.16, p = 0.88; PFC, t(14) = 1.05, p = 0.31; HIP, t(14) = 0.04, p = 0.97, (F) Kidney, t(10) = 2.30, p < 0.05; PFC, t(14) = 3.44, p < 0.01; HIP, t(14) = 0.13, p = 0.90, (G) Kidney, t(10) = 0.33, p = 0.75; PFC, t(14) = 0.74, p = 0.47; HIP, t(14) = 0.54, p = 0.60, (H) Kidney, t(10) = 3.64, p < 0.01; PFC, t(14) = 1.46, p = 0.17; HIP, t(14) = 1.28, p = 0.22, (I) Kidney, t(10) = 2.99, p < 0.05; PFC, t(14) = 1.16, p = 0.26; HIP, t(14) = 0.65, p = 0.52. HS, high salt; PFC, prefrontal cortex; HIP, hippocampus; Losartan, a non-BBB-crossing Ang II receptor blocker; PGE2, prostaglandin E2; COX-1, cyclooxygenase-1; COX-2, cyclooxygenase-2; mPGES-1, microsomal prostaglandin E synthase-1; mPGES-2, microsomal prostaglandin E synthase-2; EP1, prostaglandin E2 type 1 receptor; EP2, prostaglandin E2 type 2 receptor; EP3, prostaglandin E2 type 3 receptor; EP4, prostaglandin E2 type 4 receptor; AT8, phosphorylated tau at Ser^202^/Thr^205^ site.

The PGE2-EP1 system is reportedly involved in hypertension and neuronal damage (Cao et al., 2012, Zhen et al., 2012). Our results showed that HS intake increased EP1 mRNA levels in the kidney, as well as in the PFC (Fig 6F). Based on these results, we investigated the relationship between tau phosphorylation and the PGE2-EP1 system by co-immunostaining AT8 and EP1 using a phosphorylation-specific tau antibody in the PFC of HS-treated mice. Interestingly, tau hyperphosphorylation was merged in cells that highly expressed EP1 in the PFC of HS-treated mice (Fig 6J).

These results suggested that the PGE2-EP1 system plays an important role in tau hyperphosphorylation following HS intake.

### Hypertension, behavioral impairments, and tau hyperphosphorylation after HS intake are absent in EP1 heterozygous knockout mice

We next investigated the involvement of the PGE2-EP1 system in hypertension and behavioral impairments after HS intake in EP1^+/−^ mice. EP1^+/−^ mice were loaded with HS solution for 12 weeks and systolic blood pressure was monitored. Subsequently, locomotor activity, sociability, and object recognition memory in the mice were examined (Fig 7A). Hypertension was not observed in EP1^+/−^ mice, although EP1^+/+^ mice exhibited hypertension after HS intake (Fig 7B). Twelve weeks after HS intake, EP1+/+ mice exhibited impairments in social behavior and object recognition memory, which were not detected in EP1^+/−^ mice (Fig 7C and D). Locomotor activity and the total time taken to explore the two objects were unaltered even 12 weeks after HS intake in both EP1^+/+^ and EP1^+/−^ mice (Fig EV4B and D). To provide more direct evidence regarding the involvement of the PGE2-EP1 system in behavioral impairments, tau phosphorylation and the expression of synapse-related proteins were measured by western blotting in the PFC and HIP of EP1^+/−^ mice 12 weeks after HS intake (Fig 7E-G). Consistent with the behavioral analysis results, increased tau hyperphosphorylation was not observed in the PFC and HIP of EP1^+/−^ mice (Fig 7E). Similarly, the reduced levels of phosphorylated CaMKII in the PFC and HIP and PSD95 in the PFC were not documented in EP1^+/−^ mice (Fig 7F and G).

**Figure 7.**
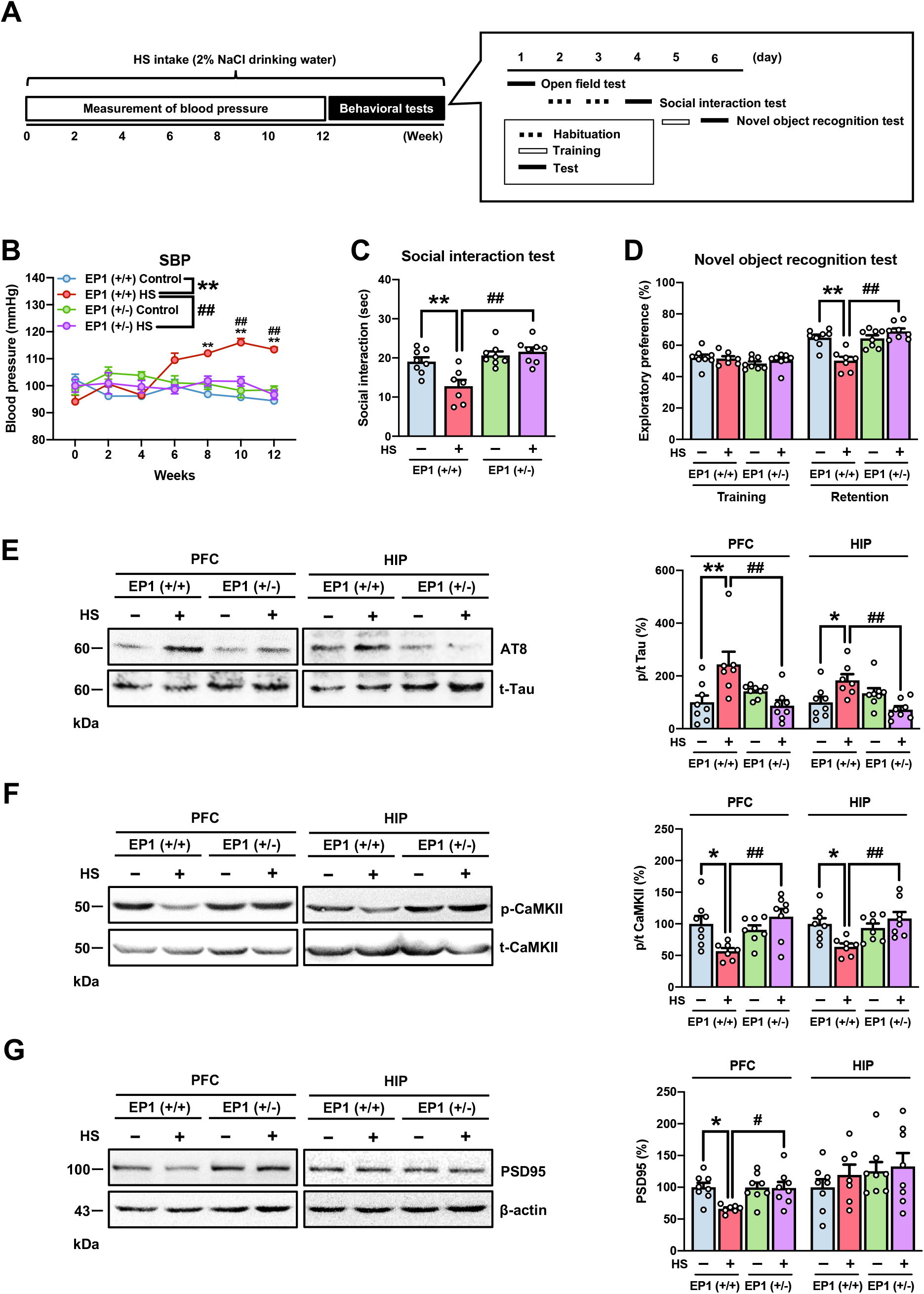
HS-induced hypertension, behavioral impairments, and tau hyperphosphorylation are not observed in heterozygous EP1 knockout mice. **A** Protocol for HS intake in heterozygous EP1 knockout mice. **B** Systolic blood pressure was monitored by the tail-cuff system every two weeks for 12 weeks. **C, D** Mice were subjected to behavioral tests 12 weeks after HS intake. (C) Social behavior in the social interaction test. The mice were placed in an apparatus with an unfamiliar mouse, and their social interaction was measured for 10 min. (D) Object recognition memory in the novel object recognition test. The retention session was performed 24 h after the training session. Exploratory preference was measured during a 10 min session. **E-G** Expression levels of phosphorylated tau and synapse-related proteins were measured in the prefrontal cortex and hippocampus of mice 12 weeks after the HS intake using western blotting. The phosphorylation ratio of (E) AT8 and (F) CaMKII and protein levels of (G) PSD95 in the prefrontal cortex and hippocampus were calculated. Loaded protein was normalized to actin. The phosphorylation ratio was calculated as tau phosphorylation versus t-Tau expression and CaMKII phosphorylation versus t-CaMKII expression. Data information: Each column represents the mean ± standard error of the mean (SEM) (n =7-8). *p < 0.05, **p < 0.01 versus EP1^+/+^ control, #p < 0.05, ##p < 0.01 versus EP1^+/+^ HS. (B)-(G) Two-way ANOVA followed by Tukey’s multiple comparison test: (B) *F* _Weeks (6,162)_ = 2.46, *p* < 0.05; *F* _group (3,27)_ = 11.76, *p* < 0.01; *F* _Weeks × group (18,162)_ = 5.41, *p* < 0.01, (C) *F* _HS (1,27)_ = 18.94, *p* < 0.01; *F* _EP1 KO (1,27)_ = 5.04, *p* < 0.05; *F* _HS × EP1 KO (1,27)_ = 9.09, *p* < 0.01, (D) Training, *F* _HS (1,27)_ = 2.24, *p* = 0.15; *F* _EP1 KO (1,27)_ = 0.17, *p* = 0.68; *F* _HS × EP1 KO (1,27)_ = 0.59, *p* = 0.45; Retention, *F* _HS (1,27)_ = 19.77, *p* < 0.01; *F* _EP1 KO (1,27)_ = 6.27, *p* < 0.05; *F* _HS × EP1 KO (1,27)_ = 21.78, *p* < 0.01, (E) PFC, *F* _HS (1,27)_ = 2.61, *p* < 0.05; *F* _EP1 KO (1,27)_ = 4.30, *p* < 0.05; *F* _HS × EP1 KO (1,27)_ = 12.73, *p* < 0.01; HIP, *F* _HS (1,27)_ = 0.31, *p* = 0.58; *F* _EP1 KO (1,27)_ = 3.97, *p* = 0.06; *F* _HS × EP1 KO (1,27)_ = 14.33, *p* < 0.01, (F) PFC, *F* _HS (1,27)_ = 1.26, *p* = 0.27; *F* _EP1 KO (1,27)_ = 5.15, *p* < 0.05; *F* _HS × EP1 KO (1,27)_ = 10.36, *p* < 0.01; HIP, *F* _HS (1,27)_ = 1.67, *p* = 0.21; *F* _EP1 KO (1,27)_ = 5.42, *p* < 0.05; *F* _HS × EP1 KO (1,27)_ = 9.81, *p* < 0.01, (G) PFC:, *F* _HS (1,27)_ = 5.50, *p* < 0.05; *F* _EP1 KO (1,27)_ = 4.85, *p* < 0.05; *F* _HS × EP1 KO (1,27)_ = 5.04, *p* < 0.05; HIP, *F* _HS (1,27)_ = 0.68, *p* = 0.42; *F* _EP1 KO (1,27)_ = 1.38, *p* = 0.25; *F* _HS × EP1 KO (1,27)_ = 0.13, *p* = 0.72. HS, high salt; SBP, systolic blood pressure; PFC, prefrontal cortex; HIP, hippocampus; AT8, phosphorylated tau at Ser^202^/Thr^205^ site; CaMKII, Ca^2+^/calmodulin-dependent protein kinase II; PSD95, postsynaptic density protein 95.

These findings strongly suggested that the PGE2-EP1 system is involved in HS-mediated hypertension and behavioral impairment.

## Discussion

It has been reported that excessive salt consumption is a risk factor for hypertension and dementia (Grillo, Salvi et al., 2019, Kendig & Morris, 2019). In the present study, we investigated the mechanisms underlying hypertension and emotional and cognitive impairments mediated by HS intake using behavioral and biochemical analyses. High dietary salt intake was found to be associated with the incidence of hypertension in human studies (Liu, 2009, Takase, Sugiura et al., 2015). Our results demonstrated that HS intake using 2% NaCl in drinking water could induce hypertension after 6 weeks, with no change in body weight or heart rate (Fig 1B, D and E). Herein, we established an HS intake model based on previous studies (Devarajan et al., 2015, Nomura et al., 2019), although HS intake was reported to elevate blood pressure after 1 to 2 weeks. The discrepancy in the onset of hypertension may be attributed to the mouse strain employed, measurement equipment, experimental procedure, or environment.

It is well-known that hypertension is one of the leading causes of chronic kidney disease (CKD) (Chen, Knicely et al., 2019). However, serum biochemical analysis, represented by BUN and Cre, suggested that hypertension 12 weeks after HS intake was not associated with renal dysfunction (Table EV1). The volume of HS (2% NaCl) consumed (Fig 1C), as well as the simultaneous levels of Na^+^ and Cl^−^ in the urine, but not serum, were markedly increased (Fig EV1D and F). Overall, these results suggest that mice may drink more to maintain electrolyte homeostasis in the blood to counteract the increased urinary excretion of Na^+^ and Cl^−^.

We performed sequential behavioral tests to evaluate changes in emotional and cognitive function at 8 and 12 weeks after HS intake. Impairments in social behavior and object recognition memory were observed after HS intake (Fig 1F and G), although no differences in anxiety-like behavior and locomotor activity were noted in the open field test, short-term memory in the Y-maze test, or spatial learning and memory in the Barnes maze test (Fig EV2A-D and F). Impaired social behavior was observed from week 8, while impaired object recognition memory was documented from week 12. Therefore, our results suggest that social behavior is more susceptible to HS intake than object recognition memory. Blood pressure was elevated 6 weeks after HS intake. Our animal model exhibits face validity, as VaD or hypertension is known to be associated with social withdrawal (Honda, Meguro et al., 2013, Redondo-Sendino, Guallar-Castillon et al., 2005). Furthermore, our results suggest that hypertension is accompanied by impairments in social behavior and object recognition memory.

It is well-established that tauopathy is pathologically defined by the deposition of abnormal tau with hyperphosphorylation and is clinically characterized by dementia (Brunden et al., 2009). Furthermore, pathological tau triggers synaptic dysfunction, which is considered a key component of the neurodegenerative process of AD (Wu et al., 2021). Our results showed hyperphosphorylation at various tau sites (Ser^202^/Thr^205^ [AT8], Ser^396^ and Ser^404^) and a reduction in synapse-related proteins (phosphorylated CaMKII and PSD95) in the PFC and HIP 12 weeks after HS intake (Fig 2B, D and E). In addition, we documented the presence of tau hyperphosphorylation 8 weeks after HS intake (Fig 2C). However, we detected no differences in the monoaminergic systems in various brain regions, including the PFC and HIP (Table EV2). Monoamine neurotransmission can be altered when a stimulus is applied (e.g., behavioral tests and psychostimulants) (Kolata, Nakao et al., 2018, Pogorelov, Kao et al., 2019). Thus, measurement of monoamines immediately after behavioral tests or the use of more temporally precise recordings, such as *in vivo* microdialysis during behavioral tests, might provide different outcomes reflecting behavioral impairments. Taken together, our results suggest that hypertension was initially induced (6 weeks after HS intake), followed by tau hyperphosphorylation, reduction of synapse-related proteins, impairment of social behavior (8 weeks intake), and impairment of recognition memory (12 weeks intake).

In addition, neuroimaging studies have shown that hypertension can be associated with reduced regional or total brain volumes (Walker, Power et al., 2017). Twelve weeks after HS intake, we observed degenerated neuronal morphology in the PrL and CA1, demonstrated as reduced dendrite length and complexity (Fig 3). The connectivity of the neuronal network, including the PFC and HIP, is known to be associated with social behavior and object recognition memory (Ko, 2017, Sorooshyari et al., 2020). In terms of cognitive function, HS intake impaired object recognition memory but not short-term memory or learning and memory. The ventral visual stream (VVS), which interacts with the PFC and HIP, plays a crucial role in visual object identification and recognition (Sorooshyari et al., 2020). Animal studies have shown that damage to the temporal cortex, one of the components of VVS, can disrupt object recognition but spare spatial memory (Bussey, Muir et al., 1999, Davies, Machin et al., 2007). Accordingly, abnormalities in neuronal morphology in the PFC and HIP following HS intake may impair the neural circuit associated with object recognition memory within the VVS.

In the RAS, Ang II, via the AT1 receptor, induces vasoconstriction and promotes aldosterone synthesis, resulting in elevated blood pressure (Sparks et al., 2014). Furthermore, the brain RAS plays an important role in oxidative stress, cerebrovascular dysfunction, and cognitive impairment (Cosarderelioglu et al., 2020, Villapol & Saavedra, 2015). Twelve weeks after HS intake, we observed that AGT and ACE mRNA levels were elevated in the kidney and/or PFC, whereas renin and AT1 mRNA levels were reduced in the HIP and PFC, respectively (Fig 4A-D). Clinical studies have shown that antihypertensive drugs, including ARBs and ACE inhibitors, can prevent dementia (Ihara & Saito, 2020). Interestingly, losartan, a non-BBB-crossing ARB, blocked hypertension, behavioral impairments, tau hyperphosphorylation, and the reduction in synapse-related proteins in the brain after HS intake (Fig 5B-G). Therefore, these findings strongly suggest that the peripheral RAS participates in HS intake-induced neuronal and behavioral impairments.

The PGE2-EP1 system is also involved in hypertension and neurotoxicity (Breyer & Breyer, 2001, Mohan et al., 2012). Epidemiological, animal, and clinical studies have revealed that nonsteroidal anti-inflammatory drugs (NSAIDs) targeting COX-1 and COX-2 attenuate the risk of AD (McGeer & McGeer, 2007). In the present study, we observed increased COX-1, COX-2, mPGES-1, EP1, EP3, and EP4 in the kidney 12 weeks after HS intake, while EP1 was increased in the PFC (Fig 6B-D, F, H and I). Interestingly, losartan blocked the increase in plasma PGE2 levels in HS-treated mice (Fig 6A). This suggests the presence of an interaction between Ang II-AT1 and PGE2-EP1 systems in HS-induced hypertension and behavioral impairments. Among PGE2 receptors, only EP1 was increased in the kidney and PFC of HS-treated mice (Fig 6F). Immunohistochemical analysis revealed that phosphorylated tau was present in cells that highly expressed EP1 and AT8 (Fig 6J). EP1 couples with Gq proteins and upregulates intracellular Ca^2+^ levels, and activation of EP1 by PGE2 induces neurotoxicity via Ca^2+^ signaling (Kawano et al., 2006). In cultured neurons, activation of EP1 by PGE2 also induces tau hyperphosphorylation (Cao, Guan et al., 2019a, Cao, Guan et al., 2019b). We found that EP1 heterozygous knockout mice did not exhibit hypertension, behavioral impairments, tau hyperphosphorylation, or reduction of synapse-related proteins in the PFC and HIP after HS intake (Fig 7B-G). Collectively, activation of the PGE2-EP1 system via Ang II-AT1 mediates HS intake-induced hypertension, as well as neuronal and behavioral impairments (Fig 8).

**Figure 8.**
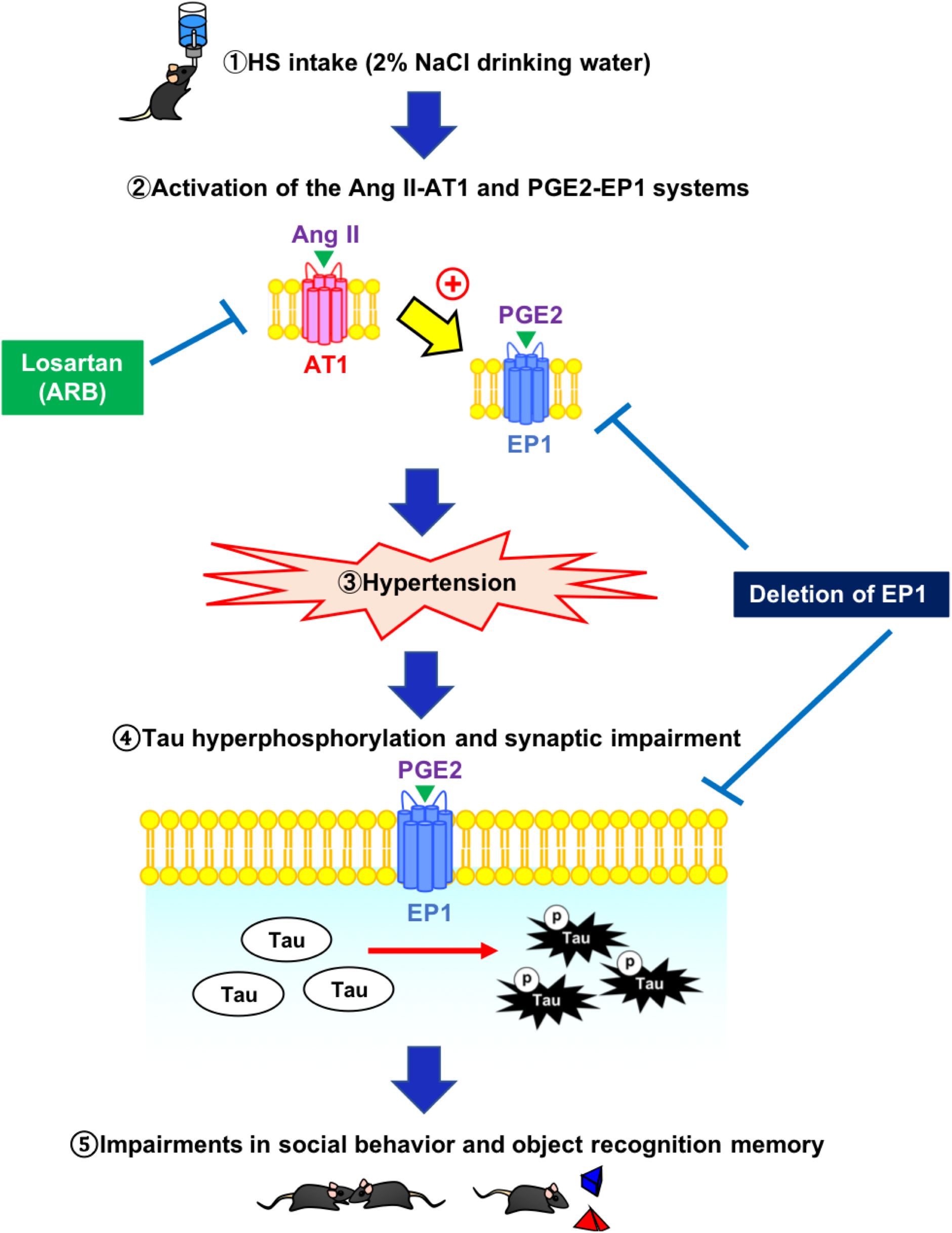
Schematic representation of the mechanism underlying HS-induced hypertension and neuronal and behavioral impairments. HS intake (①) activates the Ang II-AT1 and PGE2-EP1 systems (②), followed by hypertension (③), tau hyperphosphorylation, and reduction of synapse-related proteins in the prefrontal cortex and hippocampus (④), which are associated with impairments in social behavior and object recognition memory (⑤). Activation of the Ang II-AT1 and PGE2-EP1 systems, hypertension, and neuronal and behavioral impairments after HS intake can be blocked by losartan and the genetic knockout of EP1. HS, high salt; Ang II, angiotensin II; PGE2, prostaglandin E2; EP1, prostaglandin E2 type 1 receptor; AT1, angiotensin II type 1 receptor; losartan, a non-BBB-crossing Ang II receptor blocker.

Herein, we report a novel pathological mechanism for dementia-related to dietary HS intake. Our results demonstrate the involvement of angiotensin II-AT1 and prostaglandin E2-EP1 systems in HS intake-induced hypertension and emotional and cognitive impairments associated with tau phosphorylation. Therefore, Ang II-AT1 and PGE2-EP1 systems could be novel therapeutic targets for hypertension-induced dementia.

## Materials and methods

### Mice

In the present study, only male mice were used to exclude any potential estrous cycle effects induced by female mice. C57BL/6N mice were obtained from the Japan SLC (Shizuoka, Japan). EP1 gene-deficient mice (EP1^−/−^) were backcrossed for more than ten generations to the C57BL/6N background (Matsuoka, Furuyashiki et al., 2005). Homozygous EP1^−/−^ mice and EP1^+/+^ mice were generated by intercrossing heterozygous (EP1^+/−^) mice. We used EP1^+/−^ mice in our experiments, as EP1^−/−^ mice show impaired social behavior and impulsivity under basal conditions (Matsuoka et al., 2005). Littermate EP1^+/+^ mice were used as wild-type mice. The mice were housed in a specific pathogen-free environment within our animal facility prior to use and were maintained in a regulated environment (23 ± 3°C and 50 ± 10 % humidity) with a 12-h light/dark cycle (lights on at 8:00 A.M., off at 8:00 P.M.). Food (MF; Oriental Yeast Co., Ltd., Tokyo, Japan) and tap water were provided *ad libitum*. All experiments were performed in accordance with the guidelines established by the Japanese Pharmacological Society and Institute for Experimental Animals at Fujita Health University. The protocols were approved by the Ethics Committee of Animal Experiments of the Institute for Experimental Animals at Fujita Health University (Permit Number: AP16044).

### HS intake

An HS intake protocol was performed as previously described (Devarajan, Yahiro et al., 2015, Nomura, Hiyama et al., 2019). According to the experiment, mice (8-week-old) were forced to drink an HS solution containing 2% NaCl instead of water for 4 to 12 weeks.

### Drug administration

Losartan potassium (Tokyo Chemical Industry Co., Ltd, Tokyo, Japan) was dissolved in physiological saline and intraperitoneally (i.p.) administered daily (10:00-12:00 A.M.) during the last 6 weeks of HS intake for 12 weeks in total. The dose of losartan potassium (20 mg/kg) was selected based on a previous report (Wang, Shen et al., 2012).

### Measurement of blood pressure

Systolic blood pressure and heart rate were measured using a tail-cuff system (MK 2000; Muromachi Kikai, Tokyo, Japan), as described in a previous report (Devarajan et al., 2015) with minor modifications. Mice were acclimatized to the apparatus by performing daily measurement sessions for three days before initiating each experiment. Six blood pressure and heart rate readings were recorded each time, and the average value was used for the analysis.

### Behavioral analysis

All behavioral tests were performed between 10:00 A.M. and 6:00 P.M. Behavioral experiments were performed in a sound-attenuated and air-regulated experimental room, in which mice were habituated for more than 1 h.

The open field test was performed as described previously (Mouri, Sasaki et al., 2012) with minor modifications. The open field consisted of a square area (34 cm × 34 cm × 24 cm) set in a dark room. A light (100 lx) was positioned above the center of the apparatus floor. A mouse was placed in one corner of the open field to initiate the experiment. The mouse was then allowed to explore the open field freely. The time spent in each zone and locomotor activity were measured every 1 min for 10 min using digital counters with infrared sensors (SCANET MV-40 OF; MELQUEST Co., Ltd., Toyama, Japan).

The social interaction test was performed according to a previous report (Mouri et al., 2012) with minor modifications. The apparatus consisted of a square open arena (30 × 30 × 35 cm) with no top cover and was made of black acrylic. The light was diffused to minimize shadows in the arena and was maintained at 10 lx. A singular mouse was placed in the test box for 10 min on 2 consecutive days before the test (habituation). On the test day, each mouse was randomly assigned to a same-gender group, and age-matched C57BL/6N mice were used as an unfamiliar partner. The test mice and unfamiliar partners were placed in the test box for 10 min. The duration of social interactions (sniffing, grooming, following, mounting, and crawling, but not aggressive behavior) was recorded, and the total social interaction time was measured.

The Y-maze test was performed as previously described (Mouri et al., 2012) with minor modifications. The maze was made of gray painted wood; each arm was 40 cm long, 12 cm high, 3 cm wide at the bottom, and 10 cm wide at the top. The arms converged at an equilateral triangular central area of 4 cm along the longest axis. During an 8-min video-recorded session, each mouse was placed at the center of the apparatus and allowed to move freely through the maze. Alternation was defined as successive entries into the three arms of overlapping triplet sets. Alternation behavior (%) was calculated as the ratio of actual alternations to possible alternations, defined as (number of arm entries – 2) × 100.

The novel object recognition test was performed as previously described (Mouri et al., 2012). The test consisted of three sessions: habituation, training, and retention. Each mouse was individually habituated to a plexiglass box (30 × 30 × 35 cm high) and allowed to explore the box without any objects present for 10 min/day for 3 days (habituation session). The objects were a wooden cylinder (A), square pyramids (B), and golf balls (C), which were different in shape and color, but similar in size. During the training session, two objects (A and B) were symmetrically fixed to the floor of the box, 8 cm away from the sidewalls. A mouse was then placed midway toward the front of the box, and the total time spent exploring the two objects was recorded for a 10-min period. A mouse was considered to be exploring an object when its head was facing the object or when it was touching or sniffing the object. During the retention session, the mouse was placed back into the same box 24 h after the training session, with one familiar object (A) used during training replaced with a novel object (C). The mice were then allowed to explore freely for 10 min, and the time spent exploring each object was recorded. Throughout the experiments, the objects were counterbalanced in terms of their physical complexity and emotional neutrality. An exploratory preference index, the ratio of time spent exploring either the two objects (training session) or the novel object (retention session) to the total amount of time spent exploring both objects, was used to assess cognitive function, *e.g*., A or B/(B + A) × 100 (%) in the training session, and B or C/(B + C) × 100 (%) in the retention session.

The Barnes maze test was performed according to the method outlined in a previous report (Illouz, Madar et al., 2016). Each mouse was placed on the center of a circular table (100 cm high, 100 cm diameter, lighted 150 lx) with 20 holes (5 cm diameter) located at equal distances. Mice were allowed to explore for 120 s. Mice that could find the escape box were able to enter the box, and the test was deemed complete. Mice that failed to find the box during the allocated 120 s were guided to the box and kept in the box for 120 s. The test was performed three times daily for 5 days.

### Sample collection

Twelve weeks after HS intake, all mice were placed in metabolic cages for three days for urine collection. The mice were deeply anesthetized with isoflurane (Wako Pure Chemical Industries Co., Ltd., Osaka, Japan). The blood was collected from the abdominal inferior vena cava, and the serum was obtained by centrifugation at 800×g for 15 min at 24°C. The prefrontal cortex (PFC), hippocampus (HIP), and kidneys were rapidly dissected on ice-cold plates and frozen on dry ice. All samples were stored in a deep freezer at −80°C until use.

### Measurement of biochemical parameters

Serum levels of total protein (TP), albumin (Alb), total bilirubin (T-Bil), alkaline phosphatase (ALP), aspartate aminotransferase (AST), alanine transaminase (ALT), lactate dehydrogenase (LD), gamma-glutamyl transpeptidase (γ-GT), total cholesterol (T-cho), high-density lipoprotein cholesterol (HDL-cho), triglyceride (TG), uric acid (UA), glucose (Glu), blood urea nitrogen (BUN), and creatinine (Cre) were determined using a clinical biochemistry automated analyzer (BioMajesty JCA-BM 2250; JEOL, Tokyo, Japan).

### Electrolyte measurement

Levels of sodium (Na^+^), potassium (K^+^), and chloride (Cl^−^) in serum and urine were determined using an electrolyte analyzer, EA09 (A&T, Kanagawa, Japan).

### Measurement of PGE2

Quantification of PGE2 in plasma was performed as described previously (Eskilsson, Matsuwaki et al., 2017) with minor modifications. After 12 weeks of HS intake, blood was collected from the abdominal inferior vena cava of each mouse and transferred into EDTA-containing tubes. Indomethacin (10 μM) was immediately added to collect blood samples to prevent artificial PGE2 production, and plasma was obtained by centrifugation at 800×g for 10 min at 4°C. All samples were stored in a deep freezer at −80°C until use. PGE2 levels were determined using an enzyme immunoassay kit (Cayman Chemicals, Ann Arbor, MI, UK) according to the manufacturer’s protocol.

### Measurement of monoamines and their metabolites

Monoamines and their metabolites were measured as described previously (Mouri et al., 2012). Monoamine and metabolite contents were determined using a high-pressure liquid chromatography (HPLC) system (HTEC-500, Eicom, Kyoto, Japan). Each frozen brain sample was weighed and homogenized with an ultrasonic processor in 0.2 M perchloric acid containing isoproterenol as an internal standard. The homogenates were then placed on ice and centrifuged at 20,000×g for 15 min. The supernatants were mixed with 1 M sodium acetate to adjust the pH to 3.0 and injected into an HPLC system equipped with a reversed-phase ODS column (Eicompak SC5ODS, Eicom) and an electrochemical detector. The monoamine turnover was calculated from concentrations of monoamines and their metabolites.

### Real-time reverse transcription-PCR

The PFC, HIP, and kidney of each mouse were homogenized, and total RNA was extracted using an RNeasy Total RNA Isolation Kit (Qiagen) and converted into cDNA using a ReverTra Ace Kit (Toyobo, Osaka, Japan). Quantitative real-time PCR was performed for COX-1, COX-2, mPGES-1, and mPGES-2 using SsoFast Probe SuperMix (BioRad, Hercules, CA, USA) with the StepOne Real-Time PCR system (Life Technologies, Carlsbad, CA, USA). The primers used were COX-1 (GenBank accession number NM008969), COX-2 (GenBank accession number NM011198), mPGES-1 (GenBank accession number NM022415), mPGES-2 (GenBank accession number NM133783), and β-actin (GenBank accession number NM007393). The reaction profile consisted of an initial round at 95°C for 30 s, then 40 cycles of denaturation at 95°C for 5 s, and annealing at 60°C for 60 s in the StepOne Real-Time PCR system.

Quantitative real-time PCR was performed for AGT (angiotensinogen), renin, ACE, AT1, AT2, EP1, EP2, EP3, and EP4 using SsoAdvanced Universal SYBR Green SuperMix (Bio-Rad, Hercules, CA, USA) with a StepOne Real-Time PCR system (Life Technologies). The primers used were as follows: AGT: forward 5′-ACACCTACGTTCACTTCCAAG-3′, reverse 5′-CCGAGATGCTGTTGTCCAC-3′, Renin: forward 5′-AGGCAGTGACCCTCAACACCAG-3′, reverse 5′-CCAGTATGCAGGTCGTTCCT-3′, ACE: forward 5′-AACAAACATGATGGCCACATCCCG-3′, reverse 5′-CGTGTAGCCATTGAGCTTTGGCAAT-3′, AT1: forward 5′-CAAGTCGCACTCAAGCCTGTC-3′, reverse 5′-TCACTCCACCTCCAGAACAAGACG-3′, AT2: forward 5′-GGTCTGCTGGGATTGCCTTAATG-3′, reverse 5′-ACTTGGTCACGGGTAATTCGTTC-3′, EP1: forward 5′-CCTCGTCTGCCTCATCCATC-3′, reverse 5′-AACACCACCAACACCAGCA-3′, EP2: forward 5′-GCTCCTTGCCTTTCACAATCT-3′, reverse 5′-AGGACCGGTGGCCTAAGTAT-3′, EP3: forward 5′-TGGTCGCCGCTATTGATAATGA-3′, reverse 5′-GCAGCAGATAAACCCAGGGA-3′, EP4: forward 5′-TCATCTGCTCCATTCCGCTC-3′, reverse 5′-GGATGGGGTTCACAGAAGCA-3′ and β-actin forward 5′-CGATGCCCTGAGGCTCTTT-3′, reverse 5′-TGGATGCCACAGGATTCCA-3′. The reaction profile consisted of an initial cycle at 95°C for 30 s, then 45 cycles of denaturation at 95°C for 15 s, and annealing at 60°C for 60 s in the StepOne Real-Time PCR system.

To standardize the quantification, β-actin was simultaneously calculated with COX-1, COX-2, mPGES-1, mPGES-2, AGT, renin, ACE, AT1, AT2, EP1, EP2, EP3, and EP4. The expression levels were calculated using the ΔΔCt method.

### Immunofluorescence

Histological procedures were performed as described previously (Kunisawa, Shimizu et al., 2018) with minor modifications. Mice were anesthetized with sodium pentobarbital (70 mg/kg, i.p.) and perfused transcardially with ice-cold phosphate-buffered saline (PBS) followed by 4% paraformaldehyde in PBS. The brains were removed and post-fixed in 4% paraformaldehyde overnight at 4°C. The post-fixed tissues were cryoprotected in PBS containing 20% sucrose overnight, embedded in OCT compound (Sakura Finetechnical Co., Tokyo, Japan), and sliced into 20-μm-thick coronal sections using a cryostat (Leica CM3050, Wetzlar, Germany) for immunohistochemistry.

Cryosections were immunostained with mouse anti-phospho-tau (Ser202/Thr205) (1:200; Thermo Fisher Scientific, Waltham, MA, USA) and rabbit anti-EP1 (1:200; Cayman Chemicals). The coronal sections were heated up to 90°C in a microwave oven in 10 mM citrate buffer (pH 6.0) for 5 min. After washing with PBS containing 0.3% Triton-X (PBST), sections were blocked with 5% fetal bovine serum (Nichirei Bioscience Inc., Tokyo, Japan) in PBST for 2 h and then incubated with primary antibodies in PBST at 4°C overnight. After washing with PBST, the sections were incubated with secondary antibodies (1:2,000; Alexa488-conjugated goat anti-mouse IgG and Alexa568-conjugated goat anti-rabbit IgG; Molecular Probes, Eugene, OR, USA) and Hoechst 33342 (0.1 µg/ml; Dojindo, Kumamoto, Japan) for 3 h at room temperature. Subsequently, sections were mounted and covered with glass coverslips after rinsing with PBST and then visualized under a confocal laser microscope (Zeiss LSM-710FSX100; Olympus, Tokyo, Japan).

### Golgi staining and morphological analyses

Golgi staining was performed using the FD Rapid Golgi Stain Kit, according to the manufacturer’s protocol (FD NeuroTechnologies, Ellicott City, MD, USA) and a previous report (Wulaer, Hada et al., 2020). The mice were sacrificed by decapitation. The freshly dissected brains were immersed in solutions A and B for one week at room temperature and transferred to solution C for 24 h at 4°C. The brains were sectioned using a cryostat at a thickness of 80 μm. BZ9000 bright-field microscope (KEYENCE, Osaka, Japan) images of neurons located in the prelimbic cortex (PrL) and CA1 were obtained. Only fully impregnated neurons displaying dendritic trees without obvious truncations isolated from neighboring impregnated neurons were used for analyses. All basal dendrites within images were traced using Neurolucida software (MicroBrightField Bioscience, Williston, VT, USA) and analyzed using NeuroExplorer (MicroBrightField Bioscience). For each mouse, at least three neurons were counted for analysis.

### Western blotting analysis

Western blotting was performed as described previously (Mouri et al., 2012) with minor modifications. Tissues were homogenized in ice-cold RIPA buffer (20 mM, pH-7.4 Tris-HCl, 150 mM NaCl, 1% NP-40, 2 mM EDTA, 1mM sodium ortho-vanadate, 50 mM NaF, 0.1% sodium dodecyl sulfate (SDS), 1% sodium deoxycholate, 20 μg/ml pepstatin, 20 μg/ml aprotinin, and 20 μg/ml leupeptin) containing Complete^TM^ Mini Protease Inhibitor Cocktail (Roche Diagnostics, Mannheim, Germany) by sonication. After centrifugation at 20,000× g for 15 min at 4°C, the protein concentration in the supernatant was determined using the Bradford assay (Millipore, Billerica, MA, USA). Each protein sample was electrophoresed on 10% (w/v) SDS-polyacrylamide gel electrophoresis (PAGE) and subsequently transferred onto 0.22 μm polyvinylidene difluoride (PVDF) membranes (Millipore). Then, membranes were blocked with 5% skimmed milk in TBST for 60 min at room temperature and probed with a primary antibody at 4°C overnight. PVDF membranes were washed three times for 5 min in TBST, incubated in the appropriate HRP-conjugated secondary antibody for 2 h at room temperature, and then washed three times for 5 min using TBST. Immunoreactive bands were visualized using an ATTO LuminoGraphI (ATTO, Tokyo, Japan). The band intensities were analyzed using CS Analysis 3.0 (ATTO). The membrane was stripped in stripping buffer (100 mM 2-mercaptoethanol, 2% SDS, and 62.5 mM Tris-HCl, pH-6.7) at 55°C for 30 min.

The following primary antibodies were used: mouse anti-phospho-Tau (Ser202/Thr205) (1:1000; Thermo Fisher Scientific), rabbit anti-phospho-Tau (Ser202) (1:1,000; Cell Signaling Technology, Danvers, MA, USA), mouse anti-phospho-Tau (Ser396) (1:1,000; Cell Signaling Technology), rabbit anti-phospho-Tau (Ser404) (1:1,000; Cell Signaling Technology), rabbit anti-phospho-Tau (Ser416) (1:1,000; Cell Signaling Technology), mouse anti-Tau (1:1,000; Thermo Fisher Scientific), rabbit anti-phospho-CaMKII α (Thr286) (1:1,000; Abcam, Cambridge, UK), rabbit anti-CaMKII α (1:2,000; Sigma-Aldrich, St. Louis, MO, USA), rabbit anti-PSD95 (1:1,000; Cell Signaling Technology), and mouse anti-β-actin (1:1,000; Sigma-Aldrich). The secondary antibodies used were anti-rabbit IgG (1:2,000; Kirkegaard & Perry Laboratories, Gaithersburg, MD, USA) and anti-mouse IgG (1:2,000; Kirkegaard & Perry Laboratories).

### Statistical analysis

All data are expressed as the mean ± standard error of the mean (SEM). Statistical analyses were performed using the GraphPad Prism 8 software (GraphPad Software Inc., San Diego, CA, USA). The differences between groups were analyzed using one-way or two-way analysis of variance (ANOVA), followed by Tukey’s multiple comparison test. The Student’s t-test was used to compare two sets of data.

## Acknowledgments

We thank our lab members for their helpful discussions. This work was supported by Grants-in-Aid for Scientific Research from the Japan Society for the Promotion of Science (17H04252, 20K07931, and 20K16679) and by the Japan Science and Technology Agency (JST) FOREST Program (JPMJFR215H). In addition, this work was supported by a grant from the Education and Research Facility of Animal Models for Human Diseases at Fujita Health University, a research grant from the Smoking Research Foundation, and the Takeda Science Foundation.

## Author contributions

HK planned the project and the main conceptual ideas, conducted all experiments, and wrote the manuscript. AM supervised the study and prepared the manuscript. BW, MH, HK, and TS assisted with experiments. KK, HT, MK, SN, TN, TF, SN, and KS contributed to the discussion of the manuscript. TN supervised the study and finalized the manuscript.

---

**Figure EV1.**
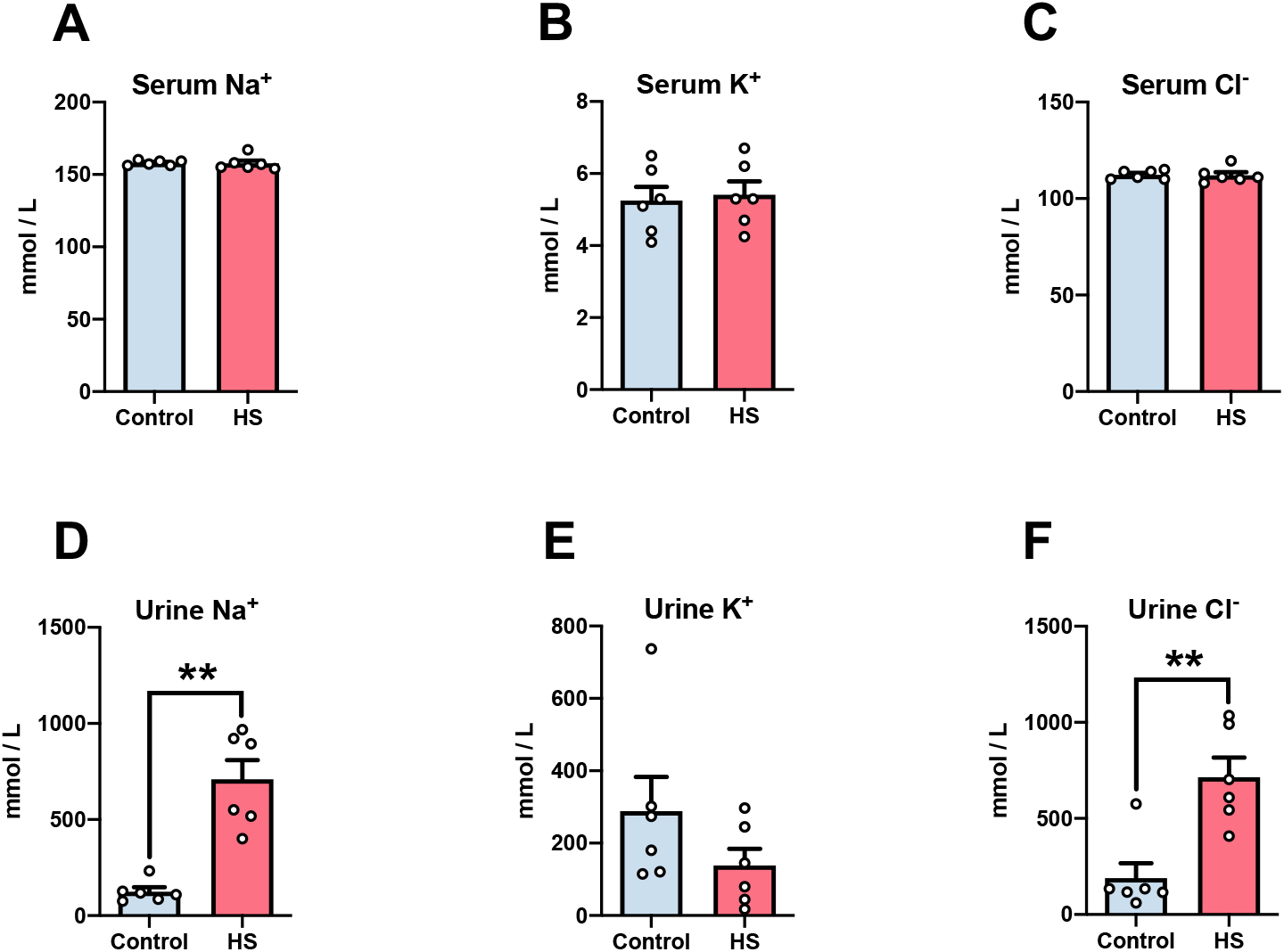
Electrolyte homeostasis after HS intake. Levels of Na^+^, K^+^, and Cl^−^ in the serum and urine of mice 12 weeks after HS intake were measured using an electrolyte analyzer, EA09. Data information: Each column represents the mean ± standard error of the mean (SEM) (n=6). **p < 0.01 versus control. (A)-(F) Student’s t-test: (A) t(10) = 0.08, p = 0.94, (B) t(10) = 0.297, p = 0.77, (C) t(10) = 0.13, p = 0.90, (D) t(10) = 5.66, p < 0.01, (E) t(10) = 1.42, p = 0.19, (F) t(10) = 4.09, p < 0.01. HS: high salt.

**Figure EV2.**
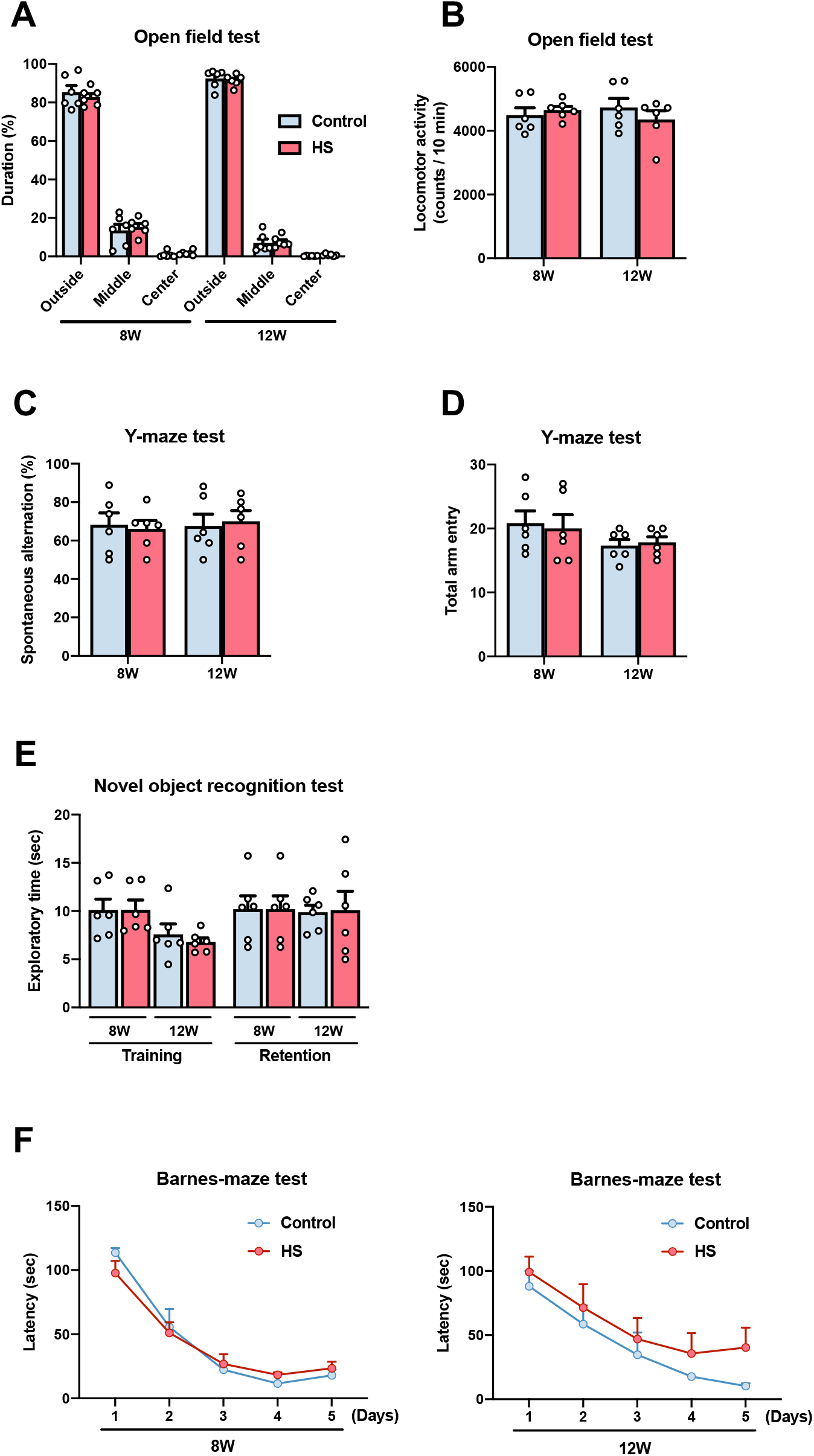
Behavioral analysis of HS-treated mice. **A-F** Mice were subjected to behavioral tests at 8 and 12 weeks after HS intake. (A), (B) Exploratory behavior in the open field test. Mice were placed in an open field, and their behavior was assessed by measuring the percentage of time spent in each zone (A) and locomotor activity (B) for 10 min. (C), (D) Short-term memory in the Y-maze test. Mice were placed in a Y-maze, and their behavior was assessed by measuring the percentage of (C) spontaneous behavior and (D) total arm entry for 8 min. (E) Exploratory time in the novel object recognition test. Exploratory time was measured during a 10 min session. (F) Spatial learning and memory in the Barnes maze test. Mice were placed at the center of the circular table, and their behavior was assessed by measuring the latency to find and enter the escape box from many holes. Data information: Each column represents the mean ± standard error of the mean (SEM) (n=6). (A), (F) Two-way ANOVA followed by Tukey’s multiple comparison test: (A) 8 weeks, *F* _Zone (1.06,10.59)_ = 533.7, *p* < 0.01; *F* _HS (1,10)_ = 0.46, *p* = 0.51; *F* _Zone × HS (2,20)_ = 0.34, *p* = 0.72; 12 weeks, *F* _Zone (1.01,10.14)_ = 1956, *p* < 0.01; *F* _HS (1,10)_ = 0.02, *p* = 0.89; *F* _Zone × HS (2,20)_ = 0.13, *p* = 0.87, (F) 8 weeks, *F* _Days (2.20,22.04)_ = 64.63, *p* < 0.01; *F* _HS (1,10)_ = 0.02, *p* = 0.89; *F* _Days × HS (4,40)_ = 1.05, *p* = 0.40; 12 weeks, *F* _Days (2.47,24.66)_ = 19.69, *p* < 0.01; *F* _HS (1,10)_ = 1.17, *p* = 0.30; *F* _Days × HS (4,40)_ = 0.36, *p* = 0.84. (B)-(E) Student’s t-test: (B) 8 weeks, t(10) = 0.62, p = 0.55; 12 weeks:, t(10) = 0.97, p = 0.36, (C) 8 weeks, t(10) = 0.29, p = 0.78; 12 weeks:, t(10) = 0.30, p = 0.77, (D) 8 weeks, t(10) = 0.29, p = 0.78; 12 weeks, t(10) = 0.39, p = 0.71, (E) Training: 8 weeks, t(10) = 0.01, p = 0.99; 12 weeks, t(10) = 0.68, p = 0.51; Retention: 8 weeks, t(10) = 0.51, p = 0.62; 12 weeks, t(10) = 0.10, p = 0.93. HS: high salt.

**Figure EV3.**
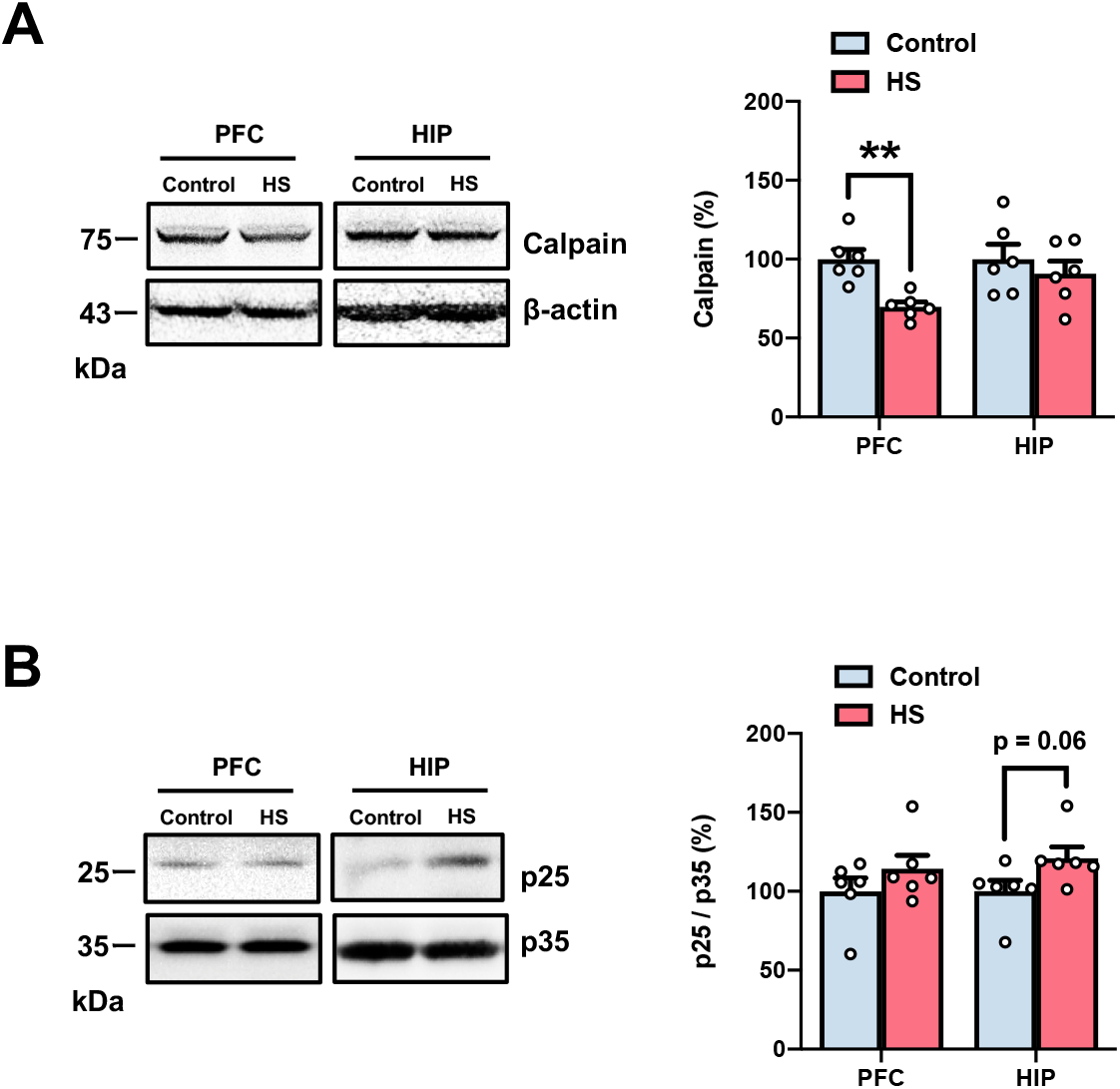
Involvement of the calpain-p25-CDK5 pathway in the prefrontal cortex and hippocampus of HS-treated mice. **A, B** Expression levels of (A) calpain and the protein ratio of (B) p25/p35 were measured by western blotting in the prefrontal cortex and hippocampus of mice 12 weeks after HS intake. Loaded protein was normalized to actin. The protein ratio was calculated as p25 versus p35. Data information: Each column represents the mean ± standard error of the mean (SEM) (n =6). **p < 0.01 versus control. (A), (B) Student’s t-test: (A) PFC, t(10) = 4.44, p < 0.01; HIP, t(10) = 0.11, p = 0.92, (B) PFC, t(10) = 1.19, p = 0.26; HIP, t(10) = 2.10, p = 0.06. HS, high salt; PFC, prefrontal cortex; HIP, hippocampus.

**Figure EV4.**
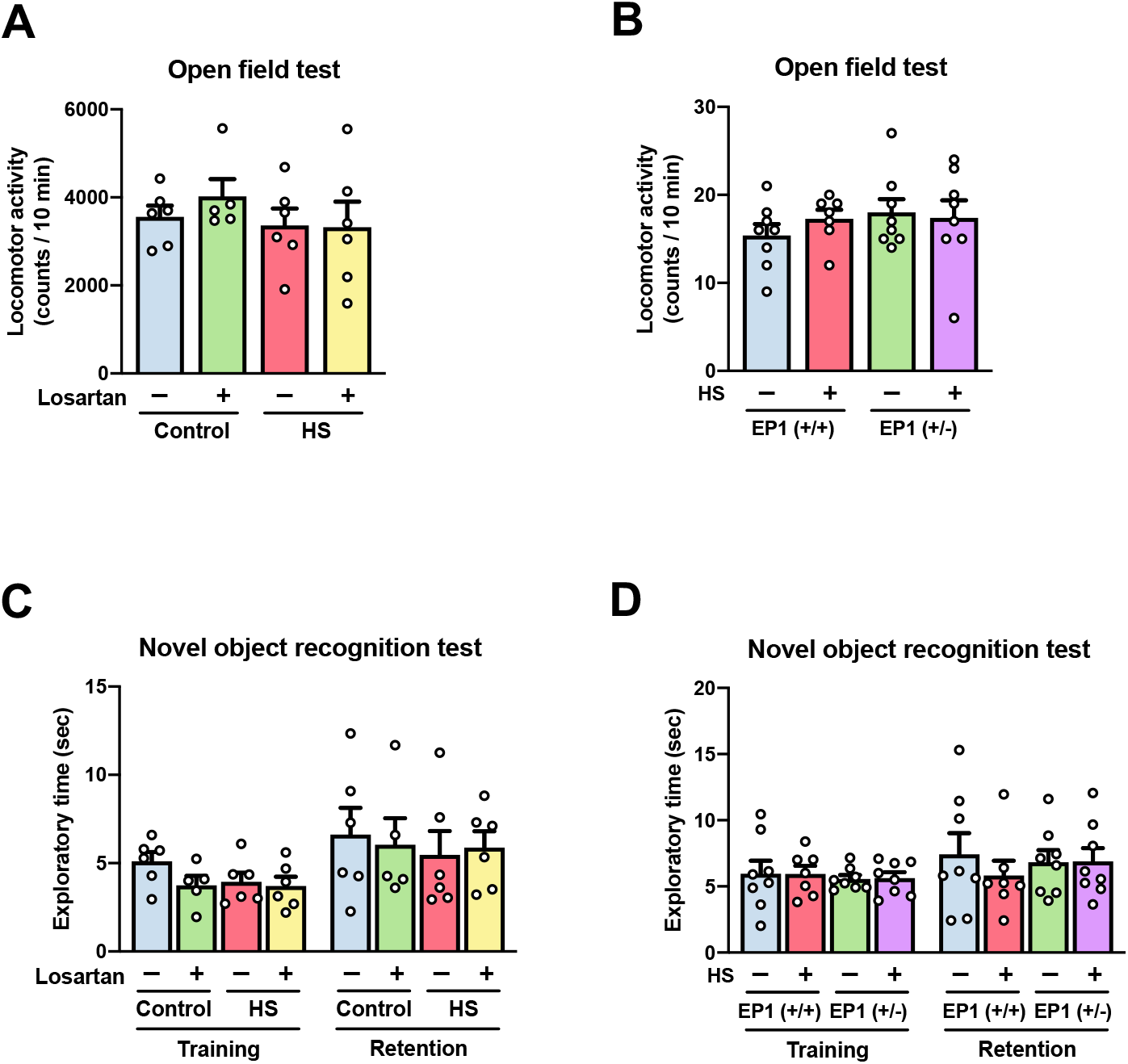
Effects of losartan and genetic EP1 knockout on object exploration and locomotor activity in the HS-treated mice. **A-D** Mice were subjected to behavioral tests 12 weeks after HS intake. (A), (B) Locomotor activity in the open field test. Mice were placed in an open field, and their locomotor activity was measured for 10 min. (C), (D) Exploratory time in the novel object recognition test. Exploratory time was measured during a 10 min session. Data information: Each column represents the mean ± standard error of the mean (SEM) (n =5-8). (A)-(D) Two-way ANOVA followed by Tukey’s multiple comparison test: (A) *F* _HS_ (1,19) = 1.11, p = 0.31; *F* _Losartan_ (1,19) = 0.25, p < 0.01; *F* _HS × Losartan_ (1,19) = 0.34, p = 0.56, (B) *F* _HS_ (1,27) = 2.15, p = 0.15; *F* _EP1 KO_ (1,27) = 0.01, p = 0.94; *F* _HS × EP1 KO_ (1,27) = 0.71, p = 0.41, (C) Training, *F* _HS_ (1,19) = 1.29, p = 0.27; *F* _Losartan_ (1,19) = 2.25, p = 0.15; *F* _HS × Losartan_ (1,19) = 1.14, p = 0.30; Retention, *F* _HS (1,19)_ = 0.15, *p* = 0.70; *F* _Losartan (1,19)_ = 1.00, *p* = 0.33; *F* _HS × Losartan (1,19)_ = 1.82, *p* = 0.19, (D) Training, *F* _HS (1,27)_ = 0.30, *p* = 0.59; *F* _EP1 KO (1,27)_ = 0.001, *p* = 0.97; *F* _HS × EP1 KO (1,27)_ = 0.004, *p* = 0.95; Retention, *F* _HS (1,27)_ = 0.04, *p* = 0.84; *F* _EP1 KO (1,27)_ = 0.42, *p* = 0.52; *F* _HS × EP1 KO (1,27)_ = 0.49, *p* = 0.49. HS, high salt; losartan, non-BBB-crossing Ang II receptor blocker.

**Table EV1.** Biochemical parameters. Biochemical parameters in the serum of mice 12 weeks after HS intake were measured using a clinical biochemistry automated analyzer (BioMajesty JCA-BM 2250). Each line represents the mean ± standard error of the mean (SEM) (n=6). HS, high salt; TP, total protein; Alb, albumin; T-Bil, total bilirubin; ALP, alkaline phosphatase; AST, aspartate aminotransferase; ALT, alanine transaminase; LD, lactate dehydrogenase; γ-GT, gamma-glutamyl transpeptidase; T-cho, total cholesterol; HDL-C, high-density lipoprotein cholesterol; TG, triglyceride; UA, uric acid; Glu, glucose; BUN, blood urea nitrogen; Cre, creatinine.

Biochemical parameters
| Serum | Control | HS |
| --- | --- | --- |
| TP (g /dl) | 5 ±0.18 | 5 ±0.12 |
| Alb (g /dl) | 0.5 ±0 | 0.58 ±0.08 |
| T-Bil (g /dl) | 0.15 ±0.03 | 0.13 ±0.01 |
| ALP (U / L) | 223 ±8.72 | 429 ±206.44 |
| AST (U / L) | 60 ±11.69 | 325 ±283.02 |
| ALT (U / L) | 36.6 ±7.49 | 205 ±185 |
| LD (U / L) | 629.1 ±55.48 | 552.5 ±151.62 |
| γ-GT (U / L) | 1.66 ±1.66 | 2.5 ±1.7 |
| T-cho (mg /dl) | 80 ±6.32 | 82.5 ±12.36 |
| HDL-cho (mg /dl) | 51.25 ±5.27 | 48.25 ±10.17 |
| TG (mg /dl) | 68.3 ±14.12 | 48.3 ±10.85 |
| UA (mg /dl) | 2.66 ±0.16 | 2.16 ±0.49 |
| Glu (mg /dl) | 127.5 ±6.02 | 140 ±21.48 |
| BUN (mg /dl) | 19.58 ±1.96 | 24.33 ±2.53 |
| Cre (mg /dl) | 0.09 ±0.01 | 0.1 ±0.01 |

**Table EV2.** Content of monoamines and their metabolites after HS intake. Levels of monoamines and their metabolites and turnovers were measured in the prefrontal cortex, nucleus accumbens, striatum, hippocampus, and amygdala of mice 12 weeks after HS intake using HPLC. Each line represents the mean ± standard error of the mean (SEM) (n=6). HS, high salt; NA, noradrenaline; MHPG, 3-methoxy-4-hydroxyphenylglycol; DA, dopamine; DOPAC, 3,4-dihydroxyphenylacetic acid; HVA, homovanillic acid; 5-HT,5-hydroxytryptamine (serotonin), 5-HIAA, 5-hydroxyindoleacetic acid.

Monoamines and their metabolites
|  | Prefrontal cortex |  | Nucleus accumbens |  | Striatum |  | Hippocampus |  | Amygdala |  |
| --- | --- | --- | --- | --- | --- | --- | --- | --- | --- | --- |
|  | Control | HS | Control | HS | Control | HS | Control | HS | Control | HS |
| NA (μmol / mg tissue) | 1.27 ± 0.07 | 1.66 ± 0.23 | 0.89 ± 0.26 | 1.11 ± 0.34 | 0.44 ± 0.08 | 0.75 ± 0.24 | 1.51 ± 0.07 | 1.44 ± 0.09 | 1.39 ± 0.08 | 1.44 ± 0.09 |
| MHPG (μmol / mg tissue) | 16.89 ± 2.54 | 15.30 ± 1.59 | 14.72 ± 1.34 | 18.52 ± 1.66 | 15.14 ± 0.74 | 16.00 ± 1.94 | 9.19 ± 0.93 | 11.18 ± 0.93 | 12.08 ± 1.66 | 14.51 ± 1.03 |
| MHPG / NA (Turnover) | 13.56 ± 2.39 | 10.06 ± 1.69 | 23.27 ± 5.77 | 40.30 ± 21.68 | 40.18 ± 6.31 | 29.62 ± 7.08 | 6.17 ± 0.67 | 7.94 ± 0.86 | 8.89 ± 1.45 | 10.35 ± 1.18 |
| DA (μmol / mg tissue) | 0.80 ± 0.19 | 5.50 ± 4.45 | 29.50 ± 2.90 | 27.06 ± 4.90 | 52.14 ± 4.60 | 50.13 ± 4.89 | 0.05 ± 0.01 | 0.10 ± 0.04 | 1.79 ± 0.07 | 1.79 ± 0.28 |
| DOPAC (μmol / mg tissue) | 0.07 ± 0.04 | 0.88 ± 0.81 | 6.07 ± 0.60 | 6.31 ± 1.81 | 12.33 ± 1.58 | 12.03 ± 2.05 | 2.13 ± 0.29 | 2.02 ± 0.19 | 0.17 ± 0.06 | 0.22 ± 0.10 |
| HVA (μmol / mg tissue) | 0.41 ± 0.09 | 1.03 ± 0.59 | 3.97 ± 0.46 | 3.72 ± 0.73 | 6.80 ± 0.40 | 7.05 ± 1.09 | 0.27 ± 0.02 | 0.33 ± 0.06 | 0.61 ± 0.04 | 0.67 ± 0.10 |
| DOPAC / DA (Turnover) | 0.09 ± 0.05 | 0.11 ± 0.03 | 0.22 ± 0.04 | 0.21 ± 0.05 | 0.24 ± 0.03 | 0.23 ± 0.03 | 42.89 ± 5.46 | 34.69 ± 9.83 | 0.10 ± 0.04 | 0.11 ± 0.05 |
| HVA / DA (Turnover) | 0.57 ± 0.09 | 0.67 ± 0.28 | 0.13 ± 0.002 | 0.14 ± 0.01 | 0.13 ± 0.01 | 0.14 ± 0.01 | 5.52 ± 0.40 | 5.15 ± 1.42 | 0.34 ± 0.02 | 0.37 ± 0.01 |
| (DOPAC+HVA) / DA (Turnover) | 0.67 ± 0.07 | 0.78 ± 0.26 | 0.36 ± 0.04 | 0.35 ± 0.05 | 0.37 ± 0.03 | 0.37 ± 0.04 | 48.41 ± 5.78 | 39.83 ± 11.19 | 0.44 ± 0.03 | 0.48 ± 0.05 |
| 5-HT (μmol / mg tissue) | 0.83 ± 0.06 | 1.08 ± 0.16 | 1.14 ± 0.07 | 1.20 ± 0.14 | 0.99 ± 0.02 | 1.08 ± 0.08 | 1.10 ± 0.06 | 1.07 ± 0.07 | 1.20 ± 0.08 | 1.19 ± 0.09 |
| 5-HIAA (μmol / mg tissue) | 0.70 ± 0.08 | 0.76 ± 0.09 | 0.72 ± 0.04 | 0.72 ± 0.10 | 1.22 ± 0.05 | 1.33 ± 0.14 | 0.75 ± 0.05 | 0.70 ± 0.12 | 0.94 ± 0.08 | 0.94 ± 0.06 |
| 5-HIAA / 5-HT (Turnover) | 0.87 ± 0.12 | 0.75 ± 0.10 | 0.63 ± 0.03 | 0.60 ± 0.06 | 1.23 ± 0.05 | 1.23 ± 0.10 | 0.69 ± 0.04 | 0.66 ± 0.12 | 0.78 ± 0.03 | 0.80 ± 0.06 |

## References

AbdAlla S, Langer A, Fu X, Quitterer U (2013) ACE inhibition with captopril retards the development of signs of neurodegeneration in an animal model of Alzheimer’s disease. Int J Mol Sci 14: 16917–42

Arvanitakis Z, Shah RC, Bennett DA (2019) Diagnosis and Management of Dementia: Review. JAMA 322: 1589–1599

Braak H, Alafuzoff I, Arzberger T, Kretzschmar H, Del Tredici K (2006) Staging of Alzheimer disease-associated neurofibrillary pathology using paraffin sections and immunocytochemistry. Acta Neuropathol 112: 389–404

Breyer MD, Breyer RM (2000) Prostaglandin E receptors and the kidney. Am J Physiol Renal Physiol 279: F12–23

Breyer MD, Breyer RM (2001) G protein-coupled prostanoid receptors and the kidney. Annu Rev Physiol 63: 579–605

Brunden KR, Trojanowski JQ, Lee VM (2009) Advances in tau-focused drug discovery for Alzheimer’s disease and related tauopathies. Nat Rev Drug Discov 8: 783–93

Bussey TJ, Muir JL, Aggleton JP (1999) Functionally dissociating aspects of event memory: the effects of combined perirhinal and postrhinal cortex lesions on object and place memory in the rat. J Neurosci 19: 495–502

Cao LL, Guan PP, Liang YY, Huang XS, Wang P (2019a) Calcium Ions Stimulate the Hyperphosphorylation of Tau by Activating Microsomal Prostaglandin E Synthase 1. Front Aging Neurosci 11: 108

Cao LL, Guan PP, Liang YY, Huang XS, Wang P (2019b) Cyclooxygenase-2 is Essential for Mediating the Effects of Calcium Ions on Stimulating Phosphorylation of Tau at the Sites of Ser 396 and Ser 404. J Alzheimers Dis 68: 1095–1111

Cao X, Peterson JR, Wang G, Anrather J, Young CN, Guruju MR, Burmeister MA, Iadecola C, Davisson RL (2012) Angiotensin II-dependent hypertension requires cyclooxygenase 1-derived prostaglandin E2 and EP1 receptor signaling in the subfornical organ of the brain. Hypertension 59: 869–76

Castro-Alvarez JF, Uribe-Arias SA, Mejia-Raigosa D, Cardona-Gomez GP (2014) Cyclin-dependent kinase 5, a node protein in diminished tauopathy: a systems biology approach. Front Aging Neurosci 6: 232

Chen TK, Knicely DH, Grams ME (2019) Chronic Kidney Disease Diagnosis and Management: A Review. JAMA 322: 1294–1304

Cohen BE, Edmondson D, Kronish IM (2015) State of the Art Review: Depression, Stress, Anxiety, and Cardiovascular Disease. Am J Hypertens 28: 1295–302

Cosarderelioglu C, Nidadavolu LS, George CJ, Marx R, Powell L, Xue QL, Tian J, Salib J, Oh E, Ferrucci L, Dincer P, Bennett DA, Walston JD, Abadir PM (2021) Higher Angiotensin II type 1 receptor (AT1R) levels and activity in the post-mortem brains of older persons with Alzheimer’s disease. J Gerontol A Biol Sci Med Sci

Cosarderelioglu C, Nidadavolu LS, George CJ, Oh ES, Bennett DA, Walston JD, Abadir PM (2020) Brain Renin-Angiotensin System at the Intersect of Physical and Cognitive Frailty. Front Neurosci 14: 586314

Davies M, Machin PE, Sanderson DJ, Pearce JM, Aggleton JP (2007) Neurotoxic lesions of the rat perirhinal and postrhinal cortices and their impact on biconditional visual discrimination tasks. Behav Brain Res 176: 274–83

Devarajan S, Yahiro E, Uehara Y, Habe S, Nishiyama A, Miura S, Saku K, Urata H (2015) Depressor effect of chymase inhibitor in mice with high salt-induced moderate hypertension. Am J Physiol Heart Circ Physiol 309: H1987–96

Disease GBD, Injury I, Prevalence C (2017) Global, regional, and national incidence, prevalence, and years lived with disability for 328 diseases and injuries for 195 countries, 1990-2016: a systematic analysis for the Global Burden of Disease Study 2016. Lancet 390: 1211–1259

Dong YF, Kataoka K, Tokutomi Y, Nako H, Nakamura T, Toyama K, Sueta D, Koibuchi N, Yamamoto E, Ogawa H, Kim-Mitsuyama S (2011) Perindopril, a centrally active angiotensin-converting enzyme inhibitor, prevents cognitive impairment in mouse models of Alzheimer’s disease. FASEB J 25: 2911–20

Eskilsson A, Matsuwaki T, Shionoya K, Mirrasekhian E, Zajdel J, Schwaninger M, Engblom D, Blomqvist A (2017) Immune-Induced Fever Is Dependent on Local But Not Generalized Prostaglandin E2 Synthesis in the Brain. J Neurosci 37: 5035–5044

Faraco G, Hochrainer K, Segarra SG, Schaeffer S, Santisteban MM, Menon A, Jiang H, Holtzman DM, Anrather J, Iadecola C (2019) Dietary salt promotes cognitive impairment through tau phosphorylation. Nature 574: 686–690

Frisoni GB, Altomare D, Thal DR, Ribaldi F, van der Kant R, Ossenkoppele R, Blennow K, Cummings J, van Duijn C, Nilsson PM, Dietrich PY, Scheltens P, Dubois B (2022) The probabilistic model of Alzheimer disease: the amyloid hypothesis revised. Nat Rev Neurosci 23: 53–66

Grillo A, Salvi L, Coruzzi P, Salvi P, Parati G (2019) Sodium Intake and Hypertension. Nutrients 11

Guan Y, Zhang Y, Wu J, Qi Z, Yang G, Dou D, Gao Y, Chen L, Zhang X, Davis LS, Wei M, Fan X, Carmosino M, Hao C, Imig JD, Breyer RM, Breyer MD (2007) Antihypertensive effects of selective prostaglandin E2 receptor subtype 1 targeting. J Clin Invest 117: 2496–505

Guo CP, Wei Z, Huang F, Qin M, Li X, Wang YM, Wang Q, Wang JZ, Liu R, Zhang B, Li HL, Wang XC (2017) High salt induced hypertension leads to cognitive defect. Oncotarget 8: 95780–95790

Honda Y, Meguro K, Meguro M, Akanuma K (2013) Social withdrawal of persons with vascular dementia associated with disturbance of basic daily activities, apathy, and impaired social judgment. Care Manag J 14: 108–13

Iadecola C (2013) The pathobiology of vascular dementia. Neuron 80: 844–66

Ihara M, Saito S (2020) Drug Repositioning for Alzheimer’s Disease: Finding Hidden Clues in Old Drugs. J Alzheimers Dis 74: 1013–1028

Illouz T, Madar R, Clague C, Griffioen KJ, Louzoun Y, Okun E (2016) Unbiased classification of spatial strategies in the Barnes maze. Bioinformatics 32: 3314–3320

Kawano T, Anrather J, Zhou P, Park L, Wang G, Frys KA, Kunz A, Cho S, Orio M, Iadecola C (2006) Prostaglandin E2 EP1 receptors: downstream effectors of COX-2 neurotoxicity. Nat Med 12: 225–9

Kehoe PG, Wong S, Al Mulhim N, Palmer LE, Miners JS (2016) Angiotensin-converting enzyme 2 is reduced in Alzheimer’s disease in association with increasing amyloid-beta and tau pathology. Alzheimers Res Ther 8: 50

Kendig MD, Morris MJ (2019) Reviewing the effects of dietary salt on cognition: mechanisms and future directions. Asia Pac J Clin Nutr 28: 6–14

Kivipelto M, Mangialasche F, Ngandu T (2018) Lifestyle interventions to prevent cognitive impairment, dementia and Alzheimer disease. Nat Rev Neurol 14: 653–666

Ko J (2017) Neuroanatomical Substrates of Rodent Social Behavior: The Medial Prefrontal Cortex and Its Projection Patterns. Front Neural Circuits 11: 41

Kolata SM, Nakao K, Jeevakumar V, Farmer-Alroth EL, Fujita Y, Bartley AF, Jiang SZ, Rompala GR, Sorge RE, Jimenez DV, Martinowich K, Mateo Y, Hashimoto K, Dobrunz LE, Nakazawa K (2018) Neuropsychiatric Phenotypes Produced by GABA Reduction in Mouse Cortex and Hippocampus. Neuropsychopharmacology 43: 1445–1456

Kunisawa K, Shimizu T, Kushima I, Aleksic B, Mori D, Osanai Y, Kobayashi K, Taylor AM, Bhat MA, Hayashi A, Baba H, Ozaki N, Ikenaka K (2018) Dysregulation of schizophrenia-related aquaporin 3 through disruption of paranode influences neuronal viability. J Neurochem 147: 395–408

Lee VM, Goedert M, Trojanowski JQ (2001) Neurodegenerative tauopathies. Annu Rev Neurosci 24: 1121–59

Liu Z (2009) Dietary sodium and the incidence of hypertension in the Chinese population: a review of nationwide surveys. Am J Hypertens 22: 929–33

Matsuoka Y, Furuyashiki T, Yamada K, Nagai T, Bito H, Tanaka Y, Kitaoka S, Ushikubi F, Nabeshima T, Narumiya S (2005) Prostaglandin E receptor EP1 controls impulsive behavior under stress. Proc Natl Acad Sci U S A 102: 16066–71

McGeer PL, McGeer EG (2007) NSAIDs and Alzheimer disease: epidemiological, animal model and clinical studies. Neurobiol Aging 28: 639–47

Milatovic D, Montine TJ, Aschner M (2011) Prostanoid signaling: dual role for prostaglandin E2 in neurotoxicity. Neurotoxicology 32: 312–9

Miners JS, Ashby E, Van Helmond Z, Chalmers KA, Palmer LE, Love S, Kehoe PG (2008) Angiotensin-converting enzyme (ACE) levels and activity in Alzheimer’s disease, and relationship of perivascular ACE-1 to cerebral amyloid angiopathy. Neuropathol Appl Neurobiol 34: 181–93

Mohan S, Ahmad AS, Glushakov AV, Chambers C, Dore S (2012) Putative role of prostaglandin receptor in intracerebral hemorrhage. Front Neurol 3: 145

Mouri A, Sasaki A, Watanabe K, Sogawa C, Kitayama S, Mamiya T, Miyamoto Y, Yamada K, Noda Y, Nabeshima T (2012) MAGE-D1 regulates expression of depression-like behavior through serotonin transporter ubiquitylation. J Neurosci 32: 4562–80

Narumiya S, Sugimoto Y, Ushikubi F (1999) Prostanoid receptors: structures, properties, and functions. Physiol Rev 79: 1193–226

Ninomiya T, Ohara T, Hirakawa Y, Yoshida D, Doi Y, Hata J, Kanba S, Iwaki T, Kiyohara Y (2011) Midlife and late-life blood pressure and dementia in Japanese elderly: the Hisayama study. Hypertension 58: 22–8

Nomura K, Hiyama TY, Sakuta H, Matsuda T, Lin CH, Kobayashi K, Kobayashi K, Kuwaki T, Takahashi K, Matsui S, Noda M (2019) [Na(+)] Increases in Body Fluids Sensed by Central Nax Induce Sympathetically Mediated Blood Pressure Elevations via H(+)-Dependent Activation of ASIC1a. Neuron 101: 60–75 e6

Oparil S, Acelajado MC, Bakris GL, Berlowitz DR, Cifkova R, Dominiczak AF, Grassi G, Jordan J, Poulter NR, Rodgers A, Whelton PK (2018) Hypertension. Nat Rev Dis Primers 4: 18014

Pogorelov VM, Kao HT, Augustine GJ, Wetsel WC (2019) Postsynaptic Mechanisms Render Syn I/II/III Mice Highly Responsive to Psychostimulants. Int J Neuropsychopharmacol 22: 453–465

Quadri SS, Culver SA, Li C, Siragy HM (2016) Interaction of the renin angiotensin and cox systems in the kidney. Front Biosci (Schol Ed) 8: 215–26

Redondo-Sendino A, Guallar-Castillon P, Banegas JR, Rodriguez-Artalejo F (2005) [Relationship between social network and hypertension in older people in Spain]. Rev Esp Cardiol 58: 1294–301

Simic G, Babic Leko M, Wray S, Harrington C, Delalle I, Jovanov-Milosevic N, Bazadona D, Buee L, de Silva R, Di Giovanni G, Wischik C, Hof PR (2016) Tau Protein Hyperphosphorylation and Aggregation in Alzheimer’s Disease and Other Tauopathies, and Possible Neuroprotective Strategies. Biomolecules 6: 6

Simic G, Babic Leko M, Wray S, Harrington CR, Delalle I, Jovanov-Milosevic N, Bazadona D, Buee L, de Silva R, Di Giovanni G, Wischik CM, Hof PR (2017) Monoaminergic neuropathology in Alzheimer’s disease. Prog Neurobiol 151: 101–138

Sorooshyari SK, Sheng H, Poor HV (2020) Object Recognition at Higher Regions of the Ventral Visual Stream via Dynamic Inference. Front Comput Neurosci 14: 46

Sparks MA, Crowley SD, Gurley SB, Mirotsou M, Coffman TM (2014) Classical Renin-Angiotensin system in kidney physiology. Compr Physiol 4: 1201–28

Takase H, Sugiura T, Kimura G, Ohte N, Dohi Y (2015) Dietary Sodium Consumption Predicts Future Blood Pressure and Incident Hypertension in the Japanese Normotensive General Population. J Am Heart Assoc 4: e001959

Takeda S, Sato N, Takeuchi D, Kurinami H, Shinohara M, Niisato K, Kano M, Ogihara T, Rakugi H, Morishita R (2009) Angiotensin receptor blocker prevented beta-amyloid-induced cognitive impairment associated with recovery of neurovascular coupling. Hypertension 54: 1345–52

Tian M, Zhu D, Xie W, Shi J (2012) Central angiotensin II-induced Alzheimer-like tau phosphorylation in normal rat brains. FEBS Lett 586: 3737–45

Ungvari Z, Toth P, Tarantini S, Prodan CI, Sorond F, Merkely B, Csiszar A (2021) Hypertension-induced cognitive impairment: from pathophysiology to public health. Nat Rev Nephrol 17: 639–654

Villapol S, Saavedra JM (2015) Neuroprotective effects of angiotensin receptor blockers. Am J Hypertens 28: 289–99

Walker KA, Power MC, Gottesman RF (2017) Defining the Relationship Between Hypertension, Cognitive Decline, and Dementia: a Review. Curr Hypertens Rep 19: 24

Wang W, Shen J, Cui Y, Jiang J, Chen S, Peng J, Wu Q (2012) Impaired sodium excretion and salt-sensitive hypertension in corin-deficient mice. Kidney Int 82: 26–33

Wu M, Zhang M, Yin X, Chen K, Hu Z, Zhou Q, Cao X, Chen Z, Liu D (2021) The role of pathological tau in synaptic dysfunction in Alzheimer’s diseases. Transl Neurodegener 10: 45

Wulaer B, Hada K, Sobue A, Itoh N, Nabeshima T, Nagai T, Yamada K (2020) Overexpression of astroglial major histocompatibility complex class I in the medial prefrontal cortex impairs visual discrimination learning in mice. Mol Brain 13: 170

Ye J, Yin Y, Liu H, Fang L, Tao X, Wei L, Zuo Y, Yin Y, Ke D, Wang JZ (2020) Tau inhibits PKA by nuclear proteasome-dependent PKAR2alpha elevation with suppressed CREB/GluA1 phosphorylation. Aging Cell 19: e13055

Yin Y, Gao D, Wang Y, Wang ZH, Wang X, Ye J, Wu D, Fang L, Pi G, Yang Y, Wang XC, Lu C, Ye K, Wang JZ (2016) Tau accumulation induces synaptic impairment and memory deficit by calcineurin-mediated inactivation of nuclear CaMKIV/CREB signaling. Proc Natl Acad Sci U S A 113: E3773–81

Young CN, Davisson RL (2015) Angiotensin-II, the Brain, and Hypertension: An Update. Hypertension 66: 920–6

Zhen G, Kim YT, Li RC, Yocum J, Kapoor N, Langer J, Dobrowolski P, Maruyama T, Narumiya S, Dore S (2012) PGE2 EP1 receptor exacerbated neurotoxicity in a mouse model of cerebral ischemia and Alzheimer’s disease. Neurobiol Aging 33: 2215–9

Zhu D, Shi J, Zhang Y, Wang B, Liu W, Chen Z, Tong Q (2011) Central angiotensin II stimulation promotes beta amyloid production in Sprague Dawley rats. PLoS One 6: e16037

